# Development of CRSIPR-Cas13a-based antimicrobials capable of sequence-specific killing of target bacteria

**DOI:** 10.1101/808741

**Authors:** Kotaro Kiga, Xin-Ee Tan, Rodrigo Ibarra-Chávez, Shinya Watanabe, Yoshifumi Aiba, Yusuke Sato’o, Feng-Yu Li, Teppei Sasahara, Bintao Cui, Moriyuki Kawauchi, Tanit Boonsiri, Kanate Thitiananpakorn, Yusuke Taki, Aa Haeruman Azam, Masato Suzuki, José R Penadés, Longzhu Cui

## Abstract

Emergence of antimicrobial-resistant bacteria is an increasingly serious threat to global health, necessitating the development of innovative antimicrobials. We established a series of CRISPR-Cas13a-based antibacterial nucleocapsid, termed CapsidCas13a(s), capable of sequence-specific killing of carbapenem-resistant *Escherichia coli* and methicillin-resistant *Staphylococcus aureus* through promiscuous RNA cleavage after recognizing corresponding antimicrobial resistance genes. CapsidCas13a constructs were generated by packaging CRISPR-Cas13a into a bacteriophage capsid to target antimicrobial resistance genes. Contrary to Cas9-based antimicrobials that lack bacterial killing capacity when the target genes are located on a plasmid, the CapsidCas13a(s) exhibited strong bacterial killing activities upon recognizing target genes regardless of their location. The antimicrobials’ treatment efficacy was confirmed using a *Galleria mellonella* larvae model. Further, we demonstrated that the CapsidCas13a(s) can assist in bacterial gene detection without employing nucleic acid amplification and optical devices.

## Main

The emergence and spread of antimicrobial resistance among pathogenic bacteria has been a growing global public health concern for several decades^1^. A recent nationwide survey of infectious disease specialists, conducted by the Emerging Infections Network of The Infectious Diseases Society of America, found that more than 60% of the participants attended to a patient with pan-resistant, untreatable bacterial infection within the previous year^2^. It is also predicted that antimicrobial-resistant (AMR) pathogens will cause 10 million fatalities per year by the year 2050 if new antimicrobial strategies are not developed^3^. In fact, the emergence of AMR bacteria has led to the present post-antibiotic era, in which many currently available antimicrobials are no longer effective. This is due in part to the decline in antibiotic innovation, as no new class of antibiotics has been developed against Gram-negative bacteria in more than 45 years, and only a limited number of antibiotics was in either phase II or III clinical trial as of 2016 [https://www.edisongroup.com/edison-explains/antibiotics/]. Therefore, there is an urgent need for new strategies to develop alternative therapeutic approaches to prevent infections of AMR bacteria.

To this end, various nucleic acid-based antibacterials, novel peptides, bacteriophage therapies, antibodies, bacteriocins, and anti-virulence compounds have been recently developed^4^. Among these, CRISPR (clustered regularly interspaced short palindromic repeats)-Cas9-encoding phages provide a means to combat such threats by selectively killing AMR bacteria. The CRISPR-Cas9 genome-editing construct, which is designed to target AMR genes, was packaged into phages to generate gene-specific antimicrobials. The resulting phage-mediated CRISPR-Cas9-based antimicrobials can be employed to selectively kill bacteria carrying targeted AMR genes^5^. However, even though the adaptability of CRISPR-Cas9 allows for the building of large libraries of CRISPR-Cas9-guided RNAs to act against rapidly evolving AMR bacteria, this strategy is ineffective against AMR bacteria carrying plasmid-encoded AMR genes, unless a lethal system, such as a toxin-antitoxin system, is also encoded by the plasmid because sole DNA cleavage does not simply result in bacterial death^6^.

CRISPR-Cas13a is the most recently identified CRISPR-Cas type VI class 2 system and is characterized by RNA-guided single-stranded RNA (ssRNA) cleavage activity^7,8^. A 2016 study by Abudayyeh *et al.* demonstrated that CRISPR-Cas13a has promiscuous ssRNA cleavage activities and slows down the host bacteria growth due to degradation of host RNAs^8^. Our group recently reported the whole genome sequencing of 11 strains of six *Leptotrichia* species and analysis of the associated CRISPR-Cas13a systems showed that *Leptotrichia shahii* Cas13a (LshCas13a) had relatively high growth inhibition effect on host cells^9^, consistent with the observation in a previous study^8^. In order to understand whether this growth inhibition was due to cell death by Cas13a-mediated cytotoxicity, we constructed two phasmid systems (pKLC21 and pKLC26, see later) carrying CRIPSR-Cas13a and target gene, respectively, and co-introduced them into *E. coli* to test their effect on the host cells. We found that LshCas13a clearly showed target sequence-specific cell killing activity if the spacer was appropriately designed (Supplementary Fig. 1). It opened a possibility that the CRISPR-Cas13a system can be applied for the development of new types of antimicrobials to cleave the mRNA of target genes regardless if the gene is encoded by the chromosome or plasmid, which might overcome the limitations of CRISPR-Cas9-based antimicrobials^6,10,11^. This possibility prompted us to explore the development of CRISPR-Cas13a-based antibacterial nucleocapsid, designated CapsidCas13a(s), which are expected to selectively kill bacteria that encode target genes on either the bacterial chromosome or plasmid. In order to determine if LshCas13a cleavage of ssRNA is sufficient to kill target bacteria, Lsh*Cas13a* and a minimal CRISPR array containing one spacer were inserted into the plasmid pC003 to generate the plasmid pKLC21, which thus was programed to target the carbapenem resistance gene *bla*_IMP-1_. The resulting pKLC21_*bla*_IMP-1_ construct was transformed into *E. coli* NEB5*α* F’l^q^ cells with or without *bla*_IMP-1_ expression using pKLC26. Growth of the *bla*_IMP-1_-expressing transformant was significantly inhibited, as compared to those without *bla*_IMP-1_ expression (Fig. 1a). Similar growth arrest was also observed in other *E. coli* cells transformed with a series of pKLC21 plasmids programed to target various carbapenem and colistin resistance genes, as well as other genes of interest (Supplementary Fig. 1). We interpreted these results as demonstrating CRISPR-Cas13a-mediated sequence-specific killing of antimicrobial-resistant *E. coli*. Following verification of the bactericidal activity of the CRISPR-Cas13a_*bla*_IMP-1_ construct against *bla*_IMP-1_-positive bacteria, AMR bacteria-specific CapsidCas13a constructs were developed by loading the CRISPR-Cas13a system onto a phage capsid. As an initial attempt, CRISPR-Cas13a_*bla*_IMP-1_ was loaded onto *E. coli* phage M13 capsid, which can infect F pili-positive *E. coli*, to generate EC-CapsidCas13a_*bla*_IMP-1_ (Fig. 1b), which demonstrated sequence-specific killing activity against bacteria carrying the *bla*_IMP-1_ gene in an EC-CapsidCas13a_*bla*_IMP-1_ concentration-dependent manner (Fig. 1c). Subsequently, the same sequence-specific killing activities were confirmed with a series of CapsidCas13a constructs programmed to target different AMR genes, including those conferring resistance to carbapenem (*bla*_IMP-1_, *bla*_OXA-48_, *bla*_VIM-2_, *bla*_NDM-1_, *bla*_KPC-2_) and colistin (*mcr*-1, *mcr*-2), all of which are the representatives of currently problematic genetic factors in the clinical setting (Fig. 1d). Conversely, non-targeting CapsidCas13a killed no bacteria under any circumstance, indicating that this construct has ideal potential for gene-directed antimicrobial therapy against AMR bacterial infections.

**Fig. 1.**
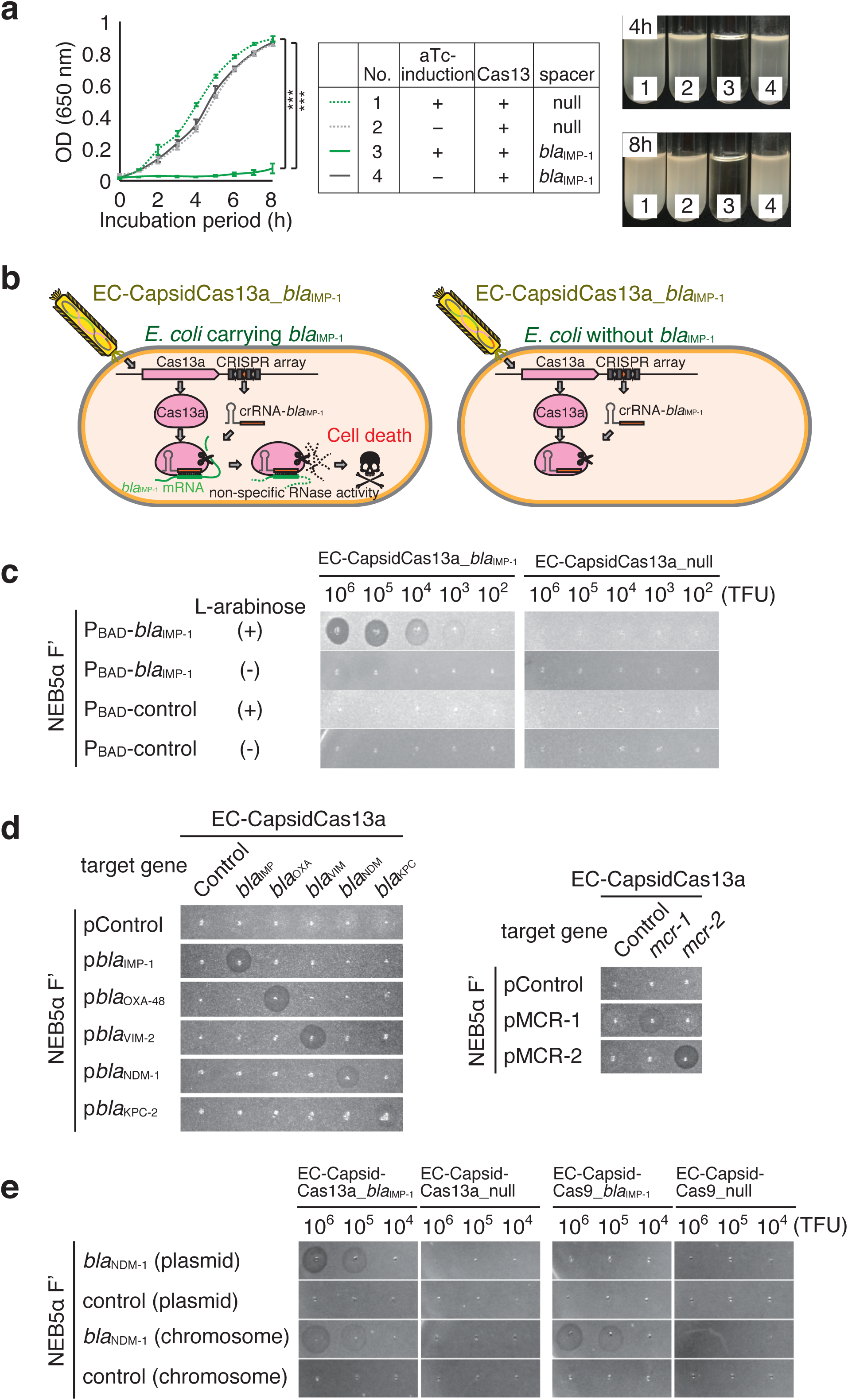
*In vitro* sequence-specific bactericidal activity of CRISPR-Cas13a from *Leptotrichia shahii* (LshCas13a). **a**, LshCas13a targeting the carbapenem resistance gene *bla*_IMP-1_ demonstrated strong killing activity against *E. coli* harboring *bla*_IMP-1_. *E. coli* DH5*α* cells were co-transformed with a CRISPR-Cas13a expression vector targeting *bla*_IMP-1_ and tetracycline-inducible *bla*_IMP-1_ expression vector with anhydrotetracycline-induced *bla*_IMP-1_ expression. Activation of the CRISPR-Cas13a system upon recognition of *bla*_IMP-1_ hindered *E. coli* growth, as shown by cell growth at an optical density at 650 nm as well as measurement of cell culture turbidity. The absence of *bla*_IMP-1_-spacer and/or the target gene *bla*_IMP-1_ did not kill the host cell, as expected, indicating that both elements are crucial for the functioning of the CRISPR-Cas13a system. **b**, Graphical concept model depicting the mode of action of CapsidCas13a. CRISPR-Cas13a_*bla*_IMP-1_ packaged onto *E. coli* M13 phage capsid (EC-CapsidCas13a_*bla*_IMP-1_) was delivered into the host bacterial cells. In the presence of the target gene (*bla*_IMP-1_-expressing *E. coli*), CRISPR-Cas13a was activated and the subsequent non-sequence-specific RNase activity resulted in host cell death. On the other hand, *E. coli* cells without the *bla*_IMP-1_ gene were unaffected. **c**, EC-CapsidCas13a-*bla*_IMP-1_ triggered cell death or growth inhibition only when the target cells expressed *bla*_IMP-1_ (L-arabinose induction). **d**, CapsidCas13a(s) programmed to target specific carbapenem resistance genes (*bla*_IMP-1_, *bla*_OXA-48_, *bla*_VIM-2_, *bla*_NDM-1_, and *bla*_KPC-2_) and colistin resistance genes (*mcr-1* and *mcr-2*) selectively killed only the bacteria harboring the corresponding resistance gene(s), as demonstrated by plaque formation. **e**, Unlike the CRISPR-Cas9 system, the CRISPR-Cas13a system selectively killed bacterial cells carrying target genes on the chromosome as well as the plasmid. Conversely, the activity of the CRISPR-Cas9 system was restricted only to the target gene located on the chromosome.

A Cas9-based EC-CapsidCas9_*bla*_NDM-1_ construct was also generated with the same protocol, but in this instance, CRISPR-Cas13a_*bla*_NDM-1_ was replaced with CRISPR-Cas9_*bla*_NDM-1_ and its bacteria-killing manner was compared with that of EC-CapsidCas13a_*bla*_NDM-1_. As expected, EC-CapsidCas9_*bla*_NDM-1_ killed only the bacteria with a chromosomal target gene, while EC-CapsidCas13a_*bla*_NDM-1_ killed the bacteria that carried the target gene on either the chromosome or plasmid (Fig. 1e). Since the majority of clinically important AMR genes are encoded on plasmids^12^, the killing efficiency of CapsidCas13a was expected to be superior to that of CapsidCas9 in light of the current difficulty with the threat of Carbapenem-resistant Enterobacteriaceae (CRE) carrying resistant genes on the plasmids. Besides, previous studies have suggested that CRISPR-Cas9 is prone to unexpected genetic mutations due to its DNA-cleavage activity^13,14,15^. On the contrary, instead of DNA, CapsidCas13a targets RNA; therefore, unintended evolution of the bacteria is unlikely.

In order to determine whether the CapsidCas13a(s) can selectively kill target bacteria among a mixed population of AMR bacteria, an artificial mixture of *E. coli* NEB5*α* F’*I*^*q*^ expressing the carbapenem resistance gene *bla*_IMP-1_, the colistin resistance gene *mcr-2*, or with no resistance gene, as a control, was treated with each gene-specific EC-CapsidCas13a construct (i.e., EC-CapsidCas13a_*bla*_IMP-1_ and EC-CapsidCas13a_*mcr-2*, respectively). Subsequently, the abundance of each bacterial strain was determined. As shown in Fig. 2a, the percentage of cell numbers of NEB5*α* F’*I*^*q*^ (*bla*_IMP-1_) and NEB5*α* F’*I*^*q*^ (*mcr-2*) were significantly decreased from 30.5% and 35% to 3.5% and 1.9%, respectively, as a result of eradicating the target cells, when the cell mixtures were treated with EC-CapsidCas13a_*bla*_IMP-1_ and EC-CapsidCas13a_*mcr-2*, respectively, while there was no change in the abundance of cell populations after treated with non-targeting EC-CapsidCas13a_null control. These highly selective killing activities of CapsidCas13a(s) demonstrate their potential as microbial control agents that can modify or manipulate the bacterial flora without affecting undesired bacterial populations by eliminating those with specific genomic contents. In addition to demonstration of the versatility of CapsidCas13a, we investigated its *in vivo* therapeutic efficacy using a *G. mellonella* larvae infection model. Administration of EC-CapsidCas13a_*bla*_IMP-1_ to *G. mellonella* larvae infected with *E. coli* R10-61 carrying *bla*_IMP-1_ showed significantly improved survival over no treatment (*p* = 0.0016) or a non-targeting CapsidCas13a_null, as a control (*p* = 0.044) (Fig. 2b). This outcome further strengthened the therapeutic prospects of the CapsidCas13a constructs for treatment of infections with AMR bacteria.

**Fig. 2.**
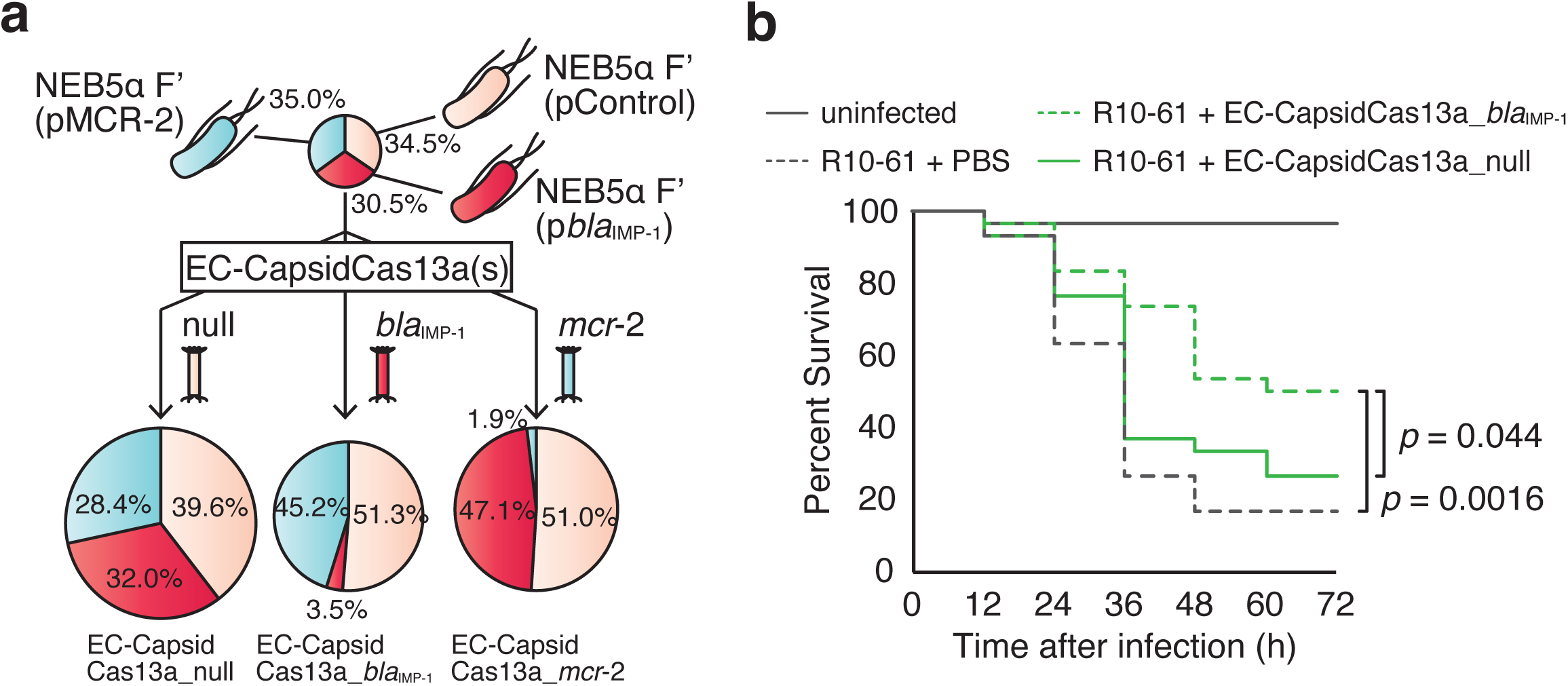
*In vivo* sequence-specific bactericidal activity of CapsidCas13a. **a**, CapsidCas13a(s) selectively eradicated the target cells from the mixed cell population. A mixed cell population was prepared by mixing *E. coli* NEB5*α* F’*I*^*q*^ with equal numbers of NEB5*α* F’*I*^*q*^ carrying *bla*_IMP-1_ and NEB5*α* F’*I*^*q*^ carrying *mcr-2*. The cells mixtures were divided into three tubes and each was treated with individual CapsidCas13a(s) targeting *bla*_IMP-1_, *mcr-2*, or non-targeting CapsidCas13a. Note that the CapsidCas13a(s) dramatically reduced the numbers of bacterial cells carrying the target genes, while the growth of non-targeted bacteria remained unchanged. Similarly, CapsidCas13a(s) with non-targeting spacer had no effect on the proportions of cells in the mixture. **b**, Time course of *G. mellonella* cell death. The therapeutic efficacies of the CapsidCas13a(s) were evaluated using a *G. mellonella* infection model. As shown by the Kaplan– Meier survival curve, administration of CapsidCas13a_*bla*_IMP-1_ targeting *bla*_IMP-1_ to the *G. mellonella* larvae infected with carbapenem-resistant clinical isolates of *E. coli* carrying *bla*_IMP-1_ significantly improved survival of the *G. mellonella* larvae, when compared to those treated with CapsidCas13a_null (control) (*p* < 0.05) or phosphate-buffered saline (*p* < 0.01) (n = 20 per group). The results are presented as the average values of three independent experiments.

Owing to the fact that the prevalence of stealth bacteria (carrying AMR genes but not identifiable by existing susceptibility tests) has been increasing^16^, the establishment of a simple and easy-to-use detection method for such strains at the genetic level is an important task in the clinical setting. Our results demonstrated that the CapsidCas13a(s) could be applied for the detection and identification of AMR genes. To this end, the CapsidCas13a constructs were modified for the detection of bacterial AMR genes. First, the spacer sequence of EC-CapsidCas13a_*bla*_IMP-1_ targeting the *bla*_IMP-1_ gene was optimized in order to improve the killing efficiency, since the bactericidal activity of CRISPR-Cas13a differed greatly depending on the spacer sequences (Supplementary Fig. 1). The optimization started with the construction of a CRISPR-Cas13a expression plasmid library, in which 121 different spacer sequences targeting *bla*_IMP-1_ were inserted into the CRISPR array. Then, all plasmids were transformed into *bla*_IMP-1_-expressing *E. coli* cells and the resulting transformants were analyzed to identify the most effective spacer sequence (Supplementary Fig. 2a). The calculation of the number of spacer reads after deep sequencing of the total plasmid DNAs extracted from the transformants found that all of the tested spacer sequences mediated target-specific bacterial killing, at least to some extent, when judged by the depletion efficiency of the plasmid DNAs (Supplementary Fig. 2b and Supplementary Table 4). Among these, 13 spacer sequences had depletion efficiencies of >100 and the *bla*_IMP-1_–563 spacer sequence GACTTTGGCCAAGCTTCTATATTTGCGT, which had the highest depletion efficiency of 364.1 was chosen as the best spacer sequence for use in subsequent experiments (Supplementary Fig. 2c). Then, the carrier M13 phage was replaced with the capsid of lysogenic phage Φ80 with the use of the PICI (phage-inducible chromosomal island) packaging system^17,18^ (Fig. 3a and Supplementary Fig. 3). The M13 phage capsid described in the previous experiment was less than ideal for clinical applications since it can only infect bacteria carrying F pili. Finally, the kanamycin resistance gene KanR was inserted as a selection marker to generate the PICI-based EC-CapsidCas13a::KanR_*bla*_IMP-1_ construct, which was expected to have improved detection sensitivity. A simulation to test the detection efficiency for *bla*_IMP-1_ was conducted by spotting 25 µL of 10-fold serial dilutions of EC-CapsidCas13a::KanR_*bla*_IMP-1_ onto fresh top agar lawns of the test strains in LB agar with or without supplementation of kanamycin. To judge the results, we opted to determine the bacterial killing effect against kanamycin-resistant cells on kanamycin plates (Fig. 3b), rather than assessing the bacterial killing effect against original cells on kanamycin-free plates (Fig. 3c). When the EC-CapsidCas13a::KanR_*bla*_IMP-1_ carrying the kanamycin-resistance gene was applied on the soft agar bacterial lawn grown on bottom agar containing kanamycin, the cells infected by this capsid acquired kanamycin resistance and, hence, could grow in the presence of kanamycin. Nevertheless, with the bacterial cells carrying the target AMR gene *bla*_IMP-1_, there was no observable growth due to the bactericidal effect of the CRISPR-Cas13a construct (Fig. 3b). This method was shown to be about 1,000 times more sensitive than direct observation of bacterial growth inhibition, as with the previous method (Fig. 1c,d and 3c). The efficiency of this system was also confirmed with the use of the M13 capsid-based EC-CapsidCas13a::KanR_*bla*_IMP-1_ construct (Supplementary Fig. 4a-c). A subsequent experiment showed that the target AMR genes of interest located on either the plasmid or chromosome could be precisely detected, demonstrating that the AMR bacteria could be identified by this method (Fig. 3d,e and Supplementary 4d,e). This finding was further confirmed by two other EC-CapsidCas13a::KanR constructs targeting *bla*_OXA-48_ and *bla*_VIM-2_ (Fig. 3d,e and Supplementary Fig. 4d,e). Although the sensitivity was slightly lower, it was even possible to detect target genes by directly spotting the CapsidCas13a(s) onto the bacteria swabbed on an agar plate instead of using the soft agar overlay method (Supplementary Fig. 5). The CapsidCas13a(s) could also simultaneously detect the toxin-encoding genes *stx*1 and *stx*2 (Fig. 3f), indicating that this method is applicable for the detection of any bacterial gene.

**Fig. 3.**
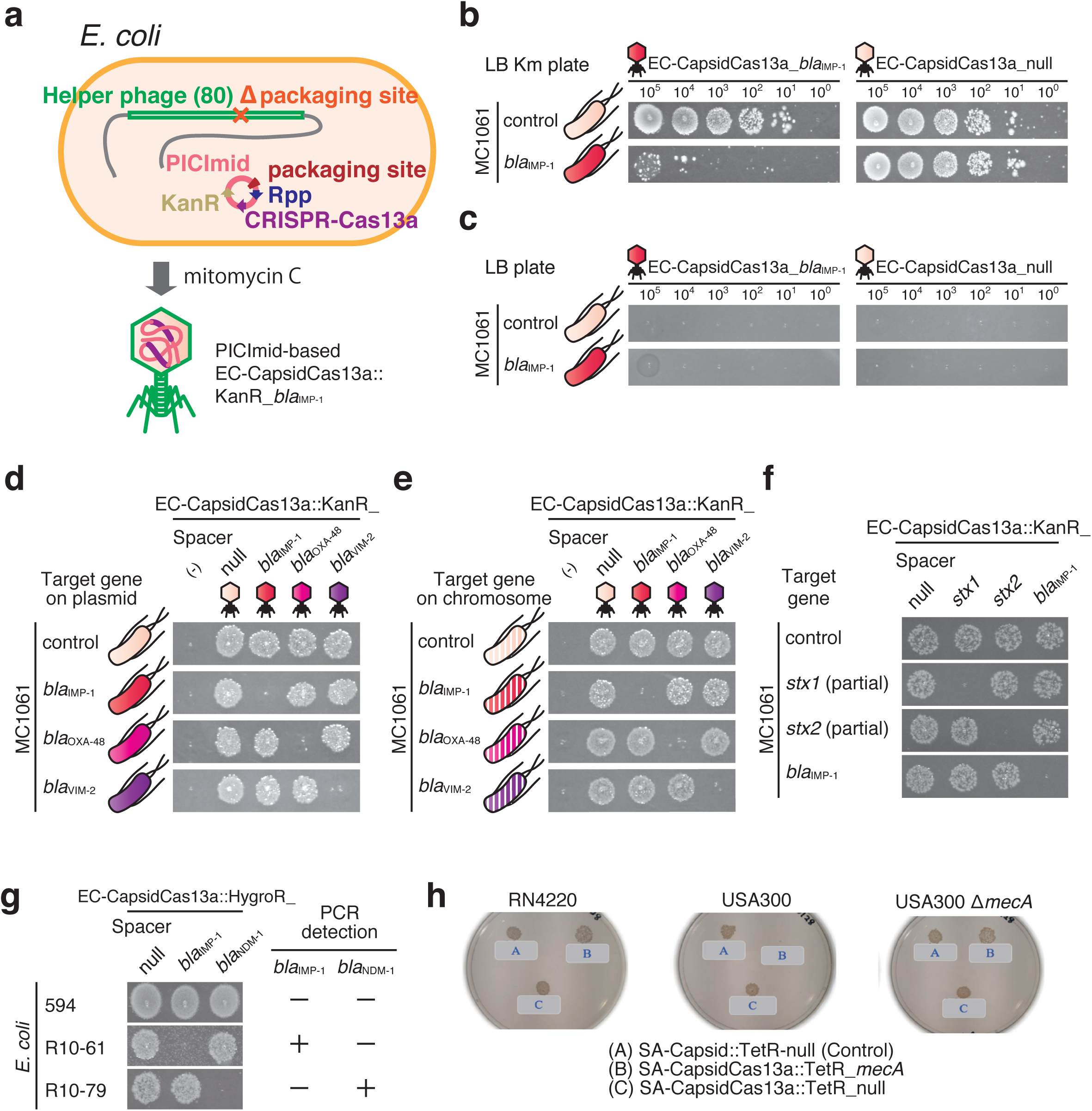
CapsidCas13a as a tool for bacterial gene detection. **a**, A schematic illustration of the generation of PICI-based EC-CapsidCas13a::KarnR_*bla*_IMP-1_ for detecting bacteria carrying *bla*_IMP-1_. **b**, In the assay for detecting *bla*_IMP-1_ gene of carbapenem-resistant *E. coli* isolates, the bacteria were mixed with soft agar and poured onto LB bottom agar plates containing kanamycin, followed by spotting of 10-fold serially diluted EC-CapsidCas13a::KanR_*bla*_IMP-1_ with the kanamycin resistance gene KanR onto the surface of the plates. **c**, The sensitivity of the test was enhanced by about 1,000 times by inserting the KanR gene into the EC-CapsidCas13a_*bla*_IMP-1_ to generate EC-CapsidCas13a::KanR_*bla*_IMP-1_ compared with the results of directly using EC-CapsidCas13a_*bla*_IMP-1_. The applicability of EC-CapsidCas13a::KanR(s) for detecting other carbapenem resistance genes (*bla*_OXA-48_, *bla*_VIM-2_, and *bla*_IMP-1_) located on both the plasmid (**d**) and chromosome (**e**) was tested. Resistance genes on either the plasmid or chromosome can be clearly detected and identified using this method. **f**, Furthermore, EC-CapsidCas13a::KanR(s) were also effectively detecting toxin-encoding genes. **g**, The carbapenem-resistant *E. coli* clinical isolate carrying *bla*_IMP-1_ was detected, where KanR was replaced with the hygromycin resistance gene HyrogR and the bottom agar contained hygromycin (200 µg/mL), as these clinical isolates were kanamycin resistant. **h**, SA-CapsidCas13a::TetR_*mecA* generated by packaging CRISPR-Cas13a targeting *mecA* of MRSA into the *S. aureus* phage capsid exhibited *mecA*-specific bacterial killing activity against MRSA but not *S. aureus* strains deficient in *mecA*.

Next, the applicability of the CapsidCas13a constructs for identification of carbapenem-resistant clinical isolates of *E. coli* carrying *bla*_IMP-1_ or *bla*_NDM-1_ was tested. As these strains were not susceptible to kanamycin, the KanR of EC-CapsidCas13a::KanR was replaced with the hygromycin resistance gene, HygroR, to generate the constructs EC-CapsidCas13a::HygroR_*bla*_IMP-1_ and EC-CapsidCas13a::HygroR_*bla*_NDM-1_. The test results with these two CapsidCas13a(s) showed that the *E. coli* isolates carrying *bla*_IMP-1_ and *bla*_NDM-1_ were clearly detected, which was consistent with the results of polymerase chain reaction (PCR) analysis (Fig. 3g). So far, there have been amazing developments of novel methods to detect bacterial genes, such as *in situ* hybridization, allele-specific PCR, loop-mediated isothermal amplification, endonuclease cleavage, direct DNA sequencing, primer extension, and oligonucleotide ligation. All these methods have unique merits in terms of sensitivity, accuracy, and robustness, allowing the wide use of each, but all have serious drawbacks that have often reduced general use. An intrinsic limitation to these molecular methods is that all are based on nucleic acid (DNA or RNA) amplification technology, which requires the use of expensive reagents as well as substantial operating costs. Reagents, disposable materials (e.g., PCR reaction tubes and pipette tips), thermal cyclers, and the necessary gel electrophoresis equipment are fairly expensive. In contrast, the proposed detection system using the CapsidCas13a constructs can be performed without amplification of DNA or RNA, electrophoresis equipment, or optical devices, as only bacterial culture plates are required. In addition, more than 10^10^ transducing forming units (TFU) of CapsidCas13a constructs can be harvested per liter of host bacterial culture, although only 2-3 µL of 10^5^ constructs per mL of solution is required for a single spot test to accurately determine the presence or absence of target genes. Together, these features highlight the elegant potentials of CapsidCas13a(s) for bacterial gene detection.

Lastly, in addition to demonstrating the bactericidal activity of CRISPR-Cas13a against Gram-negative bacteria, we also attempted to confirm the collateral activity of CRISPR-Cas13a against Gram-positive bacteria using *Staphylococcus aureus*. At first, a set of *E. coli-S. aureus* shuttle vectors, namely pKCL4(s), were generated carrying the CRISPR-Cas13a construct with or without a spacer sequence targeting the *S. aureus rpsE* genes. Transformation of the vector into *S. aureus* strain RN4220 showed that the bactericidal activity of CRISPR-Cas13a with an appropriate spacer sequence was similar to that in *E. coli* (Supplementary Fig. 6a). Then, a SA-CapsidCas13a_*mecA* construct was produced to target methicillin-resistant gene *mecA* of methicillin-resistant *S. aureus* (MRSA), one of the most prevalent AMR pathogens worldwide^19^. Optimization of the spacer sequences (Supplementary Fig. 6b-e) and the CRISPR-Cas13a_*mecA* carrying vector (pKCL-SP_*mecA*) construction were carried out in the same way as above for CRISPR-Cas13a_*bla*_IMP-1_, and packing of CRISPR-Cas13a_*mecA* into *S. aureus* phage Φ80*α* capsid was performed in accordance with the method established by Carles Ubeda *et al*. using the *S. aureus* pathogenicity island (SaPI) system^20,21^ (Supplementary Fig. 7). This packing system simultaneously imparted tetracycline resistance to the resulting SA-CapsidCas13a::TetR_*mecA* construct, since the SaPI carried the tetracycline resistance gene, which made it possible to be tested for both bacterial killing ability against MRSA and capability of MRSA detection by targeting *mecA*. As expected, when the methicillin-susceptible *S. aureus* (MSSA) strain RN4220 and MRSA USA300 or *mecA* knockout USA300 (USA300-Δ*mecA*) was infected with SA-CapsidCas13a_*mecA*, the growth of only the USA300 strain carrying *mecA* was significantly inhibited (Fig. 3h and Supplementary Fig. 8), clearly demonstrating the sequence-specific bacterial killing ability of CRISPR-Cas13a against the Gram-positive bacteria *S. aureus*.

In this study, the promiscuous RNA cleavage ability of CRISPR-Cas13a via recognition of target RNA by CRISPR-RNA (crRNA), which resulted in host cell death, was exploited to generate a new type of sequence-specific bacterial antimicrobials. To deliver the CRISPR-Cas13a to target bacteria, the CRISPR-Cas13a was packaged into carrier phage capsid using PICI packaging system. Since synthesized CapsidCas13a does not carry phage-derived genes, thereby not being able to replicate in the infected bacteria, it can be considered to be safer than genetically modified phages as being used for therapeutic. Although there are still many questions to be answered concerning the host range of the phage capsids, catalytic mode of Cas13a^22^, the efficiency of phage capsid packaging, and ethical issues regarding genetic recombinants, etc., our strategy demonstrated that CRISPR-Cas13a-based antimicrobials, CapsidCas13a(s), are likely to be developed for at least three application categories: (1) promising antibacterial therapeutic agents targeting any bacterial gene, including AMR genes, or selective killing of targeted toxin-producing bacteria; (2) a simple and inexpensive bacterial gene detection system for bacterial identification and efficient molecular epidemiological investigations without the need for the amplification of nucleic acids or optic devices; and (3) tools to manipulate the bacterial flora by targeting and eliminating a specific bacterial population without disruption to other irrelevant bacterial populations. In conclusion, the proposed CRISPR-Cas13a-based antimicrobials are expected to have a great impact in the field of antimicrobial resistance, infection control as well as bacterial flora control.

## Methods

### Bacterial strains and culture conditions

Bacterial strains used in this study are listed in Supplementary Table 1. Bacterial strains were grown at 37°C in Luria-Bertani (LB) medium (BD Difco). Unless otherwise indicated, proper antibiotics were added to growth medium to the following final concentrations: 100 µg/mL for ampicillin (Amp), 30 µg/mL for kanamycin (KM), 34 µg/mL for chloramphenicol (CP), 4 µg/mL for colistin, and 200 µg/mL for hygromycin (Hygro).

### Construction of CRISPR-Cas13a vectors targeting various genes

The CRISPR-Cas13a expression vector (pC003), which carries LshC2c2 locus on pACYC184, was kindly provided by Dr. Feng Zhang (Addgene plasmid # 79152; http://n2t.net/addgene: 79152; RRID: Addgene_79152). In order to generate an efficient vector series that carry CRISPR-Cas13a system targeting various genes, the following vector manipulations were conducted. At first, a DNA fragment of *cas13a-cas1-cas2* locus was amplified from *Leptotrichia shahii* strain JCM16776^9^ using primer set of LsCas13a clo-f and LsCas13a clo-r (Supplementary Table 3), and another DNA fragment from the pC003 vector was also amplified using primers pC003 PCR-r and pC003 PCR-f. The two DNA fragments were then ligated with In-Fusion HD Cloning kit (TaKaRa Bio Inc., Japan) to generate pKLC5, which resulted in the replacement of *cas13a-cas1-cas2* locus of pC003 with that of JCM16776. Next, the *cas1-cas2* locus was removed from the pKLC5, because Cas1/2 is not necessary for bactericidal activity of CRISPR-Cas13a. It was achieved by performing PCR on the pKLC5 using primer set of Cas1 Cas2 del SacI-f and Cas1 Cas2 del SacI-r, digesting the PCR fragment with SacI (TaKaRa Bio Inc., Japan) and ligating again with Ligation high ver. 2 (Toyobo Co., Ltd, Japan) to generate pKLC5 ΔCas1/2. Following this, in order to allow the pKLC5 ΔCas1/2 being capable to be packaged into M13 phage capsid, f1 origin together with KM resistance gene (KanR) were inserted as follows: The f1 origin and the KanR were amplified from pRC319, a gift from Timothy Lu (Addgene plasmid # 61272; http://n2t.net/addgene: 61272; RRID: Addgene_61272), using primer set of InF13 pRC319-f and InF13 pRC319-r, and another primer set of InF13 PCR SalI-f and InF13 PCR SmaI-r was used to amplify vector part from the pKLC5 ΔCas1/2 vector. The resulting two DNA fragments were then ligated with In-Fusion HD Cloning Kit to generate pKLC21 that carries complete CRISPR-Cas13a system, except for leaving the cloning site empty for insertion of appropriate spacer if necessary. The pKLC21 carries f1 origin of M13 phage thus is able to interact with M13 phage. Finally, CRISPR-Cas13a system targeting carbapenem-resistant gene *bla*_IMP-1_ was constructed on the pKLC21 vector as follows: Briefly, 33 mer oligo DNAs (28 mer of spacer and 5 mer for inserting *Bsa*I cut site) of blaIMP-1_104-s and blaIMP-1_104-as, corresponding to nucleotide position of *bla*_IMP-1_ from No. 104 to No. 132, were synthesized and annealed in annealing buffer [10 mM Tris-HCl (pH 8.0), 50 mM NaCl, 1 mM EDTA]. After that, pKLC21 was treated with restriction enzyme *Bsa*I-HF, gel-purified and subsequently ligated with the annealed oligo DNA using Ligation high ver. 2 to obtain pKLC21_*bla*_IMP-1__104. Likewise, a series of pKLC21 vectors targeting other carbapenem resistance genes (*bla*_NDM-1_, *bla*_KPC-2_, *bla*_OXA-48_, *bla*_*VIM-2*_), colistin resistance genes (*mcr*-1, *mcr*-2), as well as *mecA, rpsE*, and *ermC* of *S. aureus* were produced. The *bla*_IMP-1_-targting pKLC21 vector library covering the whole sequences of *bla*_IMP-1_ were also constructed in the same manner, whereby a total of 121 different oligo DNAs pairs covering the whole *bla*_IMP-1_ were inserted into the *Bsa*I site of pKLC21. The sequences of all oligo DNAs used were listed in the Supplementary Table 3.

Apart from pKLC21 vector series, *S. aureus*-*E. coli* shuttle vector carrying CRISPR-Cas13a (pKLC3.0) was also constructed. For the assembly of pKLC3.0, three DNA segments, KanR (*aphA-3*) and ori of *E. coli, cat* and p15A ori of *S. aureus*, and *cas13a* were amplified from plasmids pHel3, pDB114 (kindly provided by Dr. Luciano Marraffini) and pKLC21, respectively, using the following primer sets (Supplementary Table 3): InF 3.0 KanR-f and InF 3.0 KanR-r; InF 3.0 SArep_CAT-f and InF 3.0 SArep_CAT-r; and InF 3.0 p15A ori_Cas 13-f and InF 3.0 p15A ori_Cas13-r. Thereafter, the resulting 3 fragments were ligated with In-Fusion HD Cloning Kit to generate pKLC3.0. For the PICImid (plasmid carrying phage-inducible chromosomal island) construction, pBAD18 cosN *rppA*^17^ carrying three genes (c1501, c1502 and c1503) and cos site, which is necessary for the packaging of PICI of *E. coli* CFT073, was amplified with a primer set of InFpi araCosPPi NotI1-f and InFpi araCosPPi NotI1-r. The resulting fragment was digested with *Not*I (TaKaRa Bio Inc., Japan) and subsequently cloned into the *Not*I site of pKLC21 to generate PICImid pKLC31.

### Construction of *S. aureus*–*E. coli* vector containing CRISPR-Cas13a_*mecA* targeting *mecA* of MRSA

The vector pKLC-SP_*mecA* that carries CRISPR-Cas13a_*mecA* sequences flanked by two homologous recombination regions from *bap* (encoding biofilm associated protein) of Staphylococcal SaPIbov2 was constructed on the *S. aureus*-*E. coli* vector pIMAY. Two PCR fragments of 5’ and 3’ regions of *bap* on SaPIbov2 were first amplified from *S. aureus* strain RN4220-Φ80*α*-Δ*terS*-SaPIbov2::tetM^20,23^. Primers BAPup 7272-F and BAPup 8172-R were used for the PCR amplification of 5’ region of *bap* gene, while primers BAPdown 12224-F and BAPdown 13089-R were used to amplify the 3’ region. The two fragments with 15-bp flanking sequences homologous to vector ends were then sequentially integrated into a temperature-sensitive plasmid pIMAY using In-Fusion HD Cloning Kit generating pIMAY-*bap*Up/Down. Meanwhile, pKLC21_*mecA*-5 was used as a template for amplification of CRISPR-Cas13a_*mecA* with spacer targeting *mecA* and CRISPR-Cas13a system devoid of spacer targeting *mecA* (CRISPR-Cas13a_null). The PCR amplification were carried out with primer pairs of LsC2c2 mecA5-F and LsC2c2 mecA5-R to obtain CRISPR-Cas13a_*mecA*, and LsC2c2 mecA5-F and LsC2c2 188-R to obtain CRISPR-Cas13a_null (control). Finally, the PCR products were individually inserted in between 5’ and 3’ region of *bap* on pIMAY-*bap*Up/Down with In-Fusion HD Cloning Kit to generate pKLC-SP_*mecA* and pKLC-SP_null (control).

### Construction of target gene expression vectors

In this study, two plasmid systems for expression of target genes of CRISPR-Cas13a systems were constructed: pSP72 aTc-inducible vector in which the cloning site for target gene expression was under the control of anhydrotetracycline (aTc); and pKLC26 that does not contain f1 origin of replication from M13 phage. For the construction of pSP72, aTc-regulatory element was first amplified from pC008 vector using TetReg SalI-f and TetReg BamHI-r primers, and the amplicon was digested with restriction enzymes *Sal*I and *Bam*HI. After digestion, the fragment was inserted into *Sal*I-*Bam*HI site of pSP72 vector (Promega Corporation, US) carrying pBR322 origin and the resulting plasmid was termed pSP72 aTc-inducible vector because cloning site (*Bam*HI/*Eco*RI) for expression of target gene is regulated by aTc-inducible promotor. The pC008 (pBR322 with Tet-inducible RFP) was a gift from Feng Zhang (Addgene plasmid # 79157; http://n2t.net/addgene: 79157; RRID: Addgene_79157). A subsequent set of vectors expressing target genes for CRISPR-Cas13a were produced by cloning AMR genes (e.g. *bla*_IMP-1_, *bla*_NDM-1_, *bla*_KPC-2_, *bla*_OXA-48_, *bla*_*VIM-2*_) on the *Bam*HI/*Eco*RI site of the generated plasmid.

The pKLC26 was constructed basically by modifying pBAD33 with deletion of f1 origin and ARA-promoter. Firstly, pBAD33 vector was amplified with two primers, pBAD33 PCR XhoI-f and pBAD33 PCR XhoI-r, which were designed to exclude f1 origin. After digestion with restriction enzyme *Xho*I, the fragment was self-ligated with Ligation high ver. 2 to generate pKLC23. Then, the pKLC23 vector and *bla*_IMP-1_ native promoter sequence (1,716-1,168 bp of GenBank ID: AB733642.1 artificially synthesized by GENEWIZ) were independently amplified with two primer sets, pKLC23 PCR InFusion-f2 and pKLC23 PCR InFusion-r2, and InFusion-f and Intl1pro InFusion-r, respectively. Combination of the resulting two fragments with In-Fusion HD cloning kit finally generated pKLC26 vector, which carries *bla*_IMP-1_ native promoter in replacement of ARA-promoter.

To obtain pKLC26_*bla*_IMP-1_ for *bla*_IMP-1_ expression, two PCR amplifications were carried out, generating fragment of pKLC26 using a primer pair InF18 pKLC26-f and InF18 pKLC26-r, and fragment of *bla*_IMP-1_ (GenBank ID: S71932) using a primer pair InF18 IMP-1-f and InF18 IMP-1-r. The two DNA fragments were then ligated with In-Fusion HD Cloning Kit to yield pKLC26_*bla*_IMP-1_. By using the same method, a series of vectors for expression of the following AMR genes were generated (Supplementary Table 2): *bla*_NDM-1_ (GenBank ID: FN396876), *bla*_KPC-2_ (GenBank ID: AY034847), *bla*_VIM-2_ (GenBank ID: AF191564), *bla*_OXA-48_ (GenBank ID: AY236073), *mcr*-1 (GenBank ID: KP347127), *mcr*-2 (GenBank ID: LT598652) and *rfp* (encoding Red Fluorescent Protein, GenBank ID: KJ021042). The expressions of cloned genes were confirmed by susceptibility test against corresponding antibiotics using disk diffusion test recommended by CLSI.

### Generation of M13 phage-based EC-CapsidCas13a

At first, to prevent M13 phage genome from self-assembly, the f1 origin of replication of helper phage M13KO7 (New England Biolabs, US) was deleted. This was achieved by performing PCR on M13KO7 plasmid using primer pair of M13KO7 PCR InFusion-f and M13KO7 PCR InFusion-r, and subsequently ligating resulting fragment with another PCR product carrying p15A origin and CP resistance gene, which was amplified from pBAD33 using primer pair of pBAD33 PCR InFusion-f and pBAD33 PCR InFusion-r. The ligation was performed with In-Fusion HD Cloning Kit and the resulting phagemid was termed pKLC25. Next, CRISPR-Cas13a-loaded *E. coli* M13 phage capsid targeting *bla*_IMP-1_ (M13 phage-based EC-CapsidCas13a_*bla*_IMP-1_) was generated as follows: pKLC25 vector was transformed into *E. coli* MC1061 to synthesize helper phage (phage capsid). The transformed bacteria were selected on CP plates. Subsequently, the *E. coli* harboring pKLC25 was transformed with pKLC21 that carries CRISPR-Cas13a and f1 origin of M13 (i.e. pKLC21_*bla*_IMP-1_ vector) and selected on the LB agar containing CP and KM. The colonies grown on this double antibiotic selection plate were picked and cultured in LB liquid medium containing CP and KM at 37°C. *E. coli* cultures at stationary phase were then centrifuged at 8,000 *g* for 20 minutes and the supernatant was passed through a 0.22 µm filter. Equal volume of PEG buffer [5 mM Tris-HCl (pH 7.5), 10% PEG 6000, 1 M NaCl (58 g/L), 5 mM MgSO_4_·7H_2_O (1.23 g/L)] was added to the filtrate, mixed well and left at 4°C for 24 hours. After that, mixtures were centrifuged at 12,000 *g* for 10 minutes at 4°C to pellet the M13 phage-based EC-CapsidCas13a_*bla*_IMP-1_. To improve its purity, repeated centrifugation was carried out. Eventually, SM buffer [50 mM Tris-HCl (pH 7.5), 0.1 M NaCl, 7 mM MgSO_4_·7H_2_O, 0.01% gelatin] was added and the pellet was resuspended to generate a final solution of the M13 phage-based EC-CapsidCas13a_*bla*_IMP-1_. A series of M13 phage-based EC-CapsidCas13a(s) targeting the genes of *bla*_NDM-1_, *bla*_KPC-2_, *bla*_VIM-2_, *bla*_OXA-48_, *mcr*-1, *mcr*-2 and *rfp* were also prepared by using the same methods.

### Generation of PICI-based EC-CapsidCas13a and EC-CapsidCas9

The *E. coli* JP12507^18^, derived by lysogenizing phage Φ80 into *E. coli* 594 strain, was used for the generation of PICI-based EC-CapsidCas13a(s). In order to prevent Φ80 from being self-packaged during generation of PICI-based EC-CapsidCas13a, cosN site necessary for phage DNA packaging was deleted to create JP12507ΔcosN. By using JP12507ΔcosN and series of PICImid pKLC31s, PICI-based EC-CapsidCas13a(s)::KanR targeting various genes were generated as follows. At first, the pKLC31_*bla*_IMP-1_ was transformed into JP12507ΔcosN, and the transformants were cultured on KM-containing LB plates. Then, several colonies were isolated and further cultured at 37°C with shaking in LB liquid medium containing KM together with adding mitomycin C for induction of prophage excision and L-arabinose for induction of ppi (for phage DNA packaging). The L-arabinose was added to a final concentration of 0.2% that allowed to reach its final concentration of 2 µg/mL when the bacteria culture reached to OD_600_ of 0.1. The cultures were incubated overnight at 30°C with shaking at 80 rpm. After incubation, the supernatant was harvested and passed through a 0.22 µm filter. Same amount of PEG buffer was then added and the solution was left at 4°C for 1 hour after mixed well. Then, the solution was centrifuged at 12,000 *g* for 10 minutes at 4°C, resulting in precipitation of PICI-based EC-CapsidCas13a::KanR_*bla*_IMP-1_. To remove residual liquids, the pellet was washed several times and finally resuspended in SM buffer for further use. A series of PICI-based EC-CapsidCas13a::KanR(s) targeting the genes of *bla*_NDM-1_, *bla*_KPC-2_, *bla*_VIM-2_, *bla*_OXA-48_, *mcr*-1, *mcr*-2 and *rfp* were also prepared by using the same methods.

The PICI-based EC-CapsidCas13a::HygroR carrying Hygro resistance gene *hygroR* instead of KM-resistant gene *kanR* was generated by using plasmid pKLC44 in which *kanR* was placed by *hygorR*. The pKLC44 was constructed by cutting the pKLC31 with *Sac*I and *Xho*I and combining it with *hygroR* amplified from pKLC26 HygroR with primer pair of InF44 pKLC42-f and InF44 pKLC42-r using In-Fusion HD Cloning Kit.

PICI-based EC-CapsidCas9 targeting *bla*_NDM-1_ (EC-CapsidCas9_*bla*_NDM-1_) was generated using JP12507ΔcosN strain and plasmid pKLC27 (carrying CRISPR-Cas9_*bla*_NDM-1_ and packaging site cosN *rppA*. The pKLC27 was constructed as follows. The CRISPR-Cas9_*bla*_NDM-1_ was amplified from the pCR319 (a gift from Timothy Lu)^6^ using primer set of InF27 pRC319-f and InF27 pRC319-r, and packaging site cosN *rppA* was amplified from pBAD18 cosN *rppA* using primers of InF27 araCosPPi-f and InF27 araCosPPi-r. The two amplicons were then ligated with In-Fusion HD Cloning Kit to generate pKLC27.

### Generation of SaPI-based SA-CapsidCas13a_*mecA*

The SaPI-based SA-CapsidCas13a_*mecA* was generated basically using SaPI (*Staphylococcus aureus* pathogenicity island) packaging system^11,21^. At first, the *S. aureus-E.coli* shuttle vector pKLC-SP_*mecA* carrying CRISPR-Cas13a_*mecA* targeting *mecA* of MRSA and control vector pKLC-SP_null were individually transformed into *S. aureus* strain RN4220-Φ80*α*-Δ*terS*-SaPIbov2::tetM^20,23^ by electroporation using NEPA21 electroporator (Nepa Gene Co., Ltd., Japan) with the following parameters: Poring Pulse (Voltage: 1,800 V, Pulse length: 2.5 msec, Pulse interval: 50 msec, Number of pulse: 1, Polarity: +), Transfer Pulse (Voltage: 100 V, Pulse length: 99 msec, Pulse interval: 50 msec, Number of pulse: 5, Polarity: ±). The resulting transformants were then recovered at a temperature permissive for the plasmid replication (28°C) for 1 hour and plated on tryptic soy agar (TSA) plates supplemented with 10 µg/mL CP. The plasmid can be integrated into the chromosome by homologous recombination when the transformants were incubated at 37°C (non-permissive temperature) in the presence of CP. Single colonies (from 37°C plate) were then inoculated into tryptic soy broth (TSB) and incubated at 28°C without antibiotic selection to stimulate rolling cycle replication. Finally, 100 µl of the 10^−5^ dilution of this culture is plated on TSA with 1 µg/mL anhydrotetracycline to select for cells free of plasmid. Insertion of both CRISPR-Cas13a systems are validated by PCR. Finally, resulting cells were chemically induced by mitomycin C to generate SA-CapsidCas13a_*mecA* and SA-CapsidCas13a_null (control) by using the method described elsewhere^11^.

### Generation of *E. coli* strain carrying foreign gene on its own chromosome

Generation of *E. coli* strain that express foreign gene on its own chromosome was carried out by using ARA-inducible Red recombination system. To this end, at first, we transformed *E. coli* strains NEB 5-alpha F’l^q^ (New England Biolabs, US) and MC1061 with plasmid pKD46 which carries ARA-inducible Red recombination system^24^. Next, the desired genes, e.g. *bla*_IMP-1_, were knocked-in into above strains following the methods described by Tomoya Baba *et al*^25^. Briefly, a DNA fragment containing *bla*_IMP-1_ and CP resistance gene (*cat*) was amplified from the pKLC26_*bla*_IMP-1_ with primer pair of K12 genome-in pKLC26-f and K12 genome-in pKLC26 Cm-r, and the PCR product was electroporated into the above *E. coli* cell using ELEPO 21 with the following parameters: Poring Pulse (Voltage: 2,000 V, Pulse length: 2.5 msec, Pulse interval: 50 msec, Number of pulses: 1, Polarity: +), Transfer Pulse (Voltage: 150 V, Pulse length: 50 msec, Pulse interval: 50 msec, Number of pulses: 5, Polarity: ±). After electroporation, 1 mL of SOC was added and the mixture was cultured at 37°C for 1 hour with agitation, before spreading on LB plate containing CP and further incubated at 37°C until colonies were observed. The sequence of the resulting insertion was confirmed by Sanger sequencing.

### Sequence-specific bacterial killing by EC-CapsidCas13a

The logarithmic phase cultures of three *E. coli* strains with overexpression of *bla*_IMP-1_, *mcr-2* in plasmid and carrying control plasmid, NEB 5-alpha F’l^q^ (pKLC26_*bla*_IMP-1_), and NEB 5-alpha F’l^q^ (pKLC26_*mcr-2*) and NEB 5-alpha F’l^q^ (pKLC26), in LB liquid medium were adjusted to an OD_600_ of 0.1, and diluted 10 times with fresh LB medium. Transferred 300*μ*l of each these dilutions into one tube, mixed well, then divided them into four tubes up to 100*μ*l for each tube. Then, three tubes were treated with M13-based EC-CapsidCas13a_*bla*_IMP-1_, EC-CapsidCas13a_*mcr-2*, and EC-CapsidCas13a_null (control), respectively, the remaining one tube was regarded as non-treated control. EC-CapsidCa13a treatments were carried out by individually adding 100 MOI of the above three CapsidCas13a(s) to their corresponding tubes, mixed well, then culturing at 37°C with gentle shaking for 6 hours. After incubation, 10-fold serial dilutions were made with 0.8% NaCl and colony count for calculating number of survived cells was carried out. The number of colonies formed on the agar plates containing Amp, colistin and drug-free, respectively, were counted, and cell ratios were calculated.

Sequence-specific killing activity of SA-CapsidCas13a_*mecA* was determined by using tetracycline (Tet) agar plate, since SA-CapsidCas13a(s) generated were carrying Tet resistance determinant delivered from SaPIbov2 during the packaging processes.

### Measurement of phage titres (Transduced colony-forming units, TFU/mL)

The M13 phage/PICI-based EC-CapsidCas13a(s) were serially 10-fold diluted with SM buffer ranging from 10^−4^ to 10^−7^. In the meantime, overnight culture of *E. coli* strain NEB 5-alpha F’l^q^ or MC1061 diluted 1: 100 with LB broth was incubated with agitation at 37°C until an OD_600_ of approximately 0.1. Then, 10 µl of each dilution of M13 phage/PICI-based EC-CapsidCas13a(s) was added to 100 µl of bacterial suspension and the mixture was incubated at 37°C for 30 minutes. Subsequently, all of the culture solution was plated on LB plates containing CP or KM, and the plates were incubated overnight at 37°C. The colonies grown on the KM plate but not on the CP were counted to calculate the TFU/mL.

### Detection of bacterial genes with CapsidCas13a(s)

The *E. coli* strains to be determined were grown to an OD_600_ of about 0.5. Then, 100 µl of the culture were mixed with 3 mL molten soft agar (LB solution with 0.5% agarose) prewarmed at 50°C and pour onto an LB plate containing KM or Hygro. The plates were solidified at room temperature. Meanwhile, M13 phage/PICI-based EC-CapsidCas13a with known titres were adjusted to 10^5^ TFU/mL and its 10-fold serial dilutions were prepared. Finally, 2 µL of each dilution of the M13 phage/PICI-based EC-CapsidCas13a were spotted onto the solidified soft agar and the plates were incubated at 37°C. The result was interpreted as positive if bactericidal plaque was formed on the plate.

### pKLC21_*bla*_IMP-1_ library sequencing

The pKLC21_*bla*_IMP-1_ (CRISPR-Cas13a_*bla*_IMP-1_ expression plasmid) library targeting the whole region of *bla*_IMP-1_ were constructed as aforementioned. Equal amount of each 121 pKLC21_*bla*_IMP-1_ carrying spacers against different position of *bla*_IMP-1_ and pKLC21_null (as a control) were mixed and transformed into both *E. coli* MC1061(pKLC26) and MC1061(pKLC26_*bla*_IMP-1_). Each transformant was then plated onto 20 LB plates containing both CP and KM, and incubated at 37°C for 16 hours. After that, a total of more than 10,000 colonies for each transformant were harvested by using LB medium, and plasmids were extracted using QIAGEN Plasmid Midi Kit (QIAGEN). Plasmid sequencing was performed using Illumina MiSeq platform (2 x 301 bp) as described previously^9^.

### *Galleria mellonella* survival assay

The M-sized *Galleria mellonella* larvae purchased from Ikiesa company (Osaka, Japan) were used for the survival assay to assess the effect of PICI-based EC-CapsidCas13a on treatment of *E. coli* infections. Upon receipt, the larvae were acclimated to the laboratory environment by leaving them in the dark room for 24 hours before starting the assay. Larvae with weak movement, dark color, unusual shape, and sizes that differ distinctly from other larvae were excluded from the experiment. Hamilton syringe (701LT, Hamilton) and a KF 731 needle (Hamilton) were used in this experiment. At first, overnight culture of carbapenem-resistant *E. coli* R10-16 carrying carbapenem resistance gene *bla*_IMP-1_ was diluted at 1: 1000 with fresh LB medium and further incubated at 37°C with agitation to reach an OD_600_ of about 0.5. The bacteria were then washed twice with PBS and adjusted to a density of approximately 1 × 10^7^ CFU/mL in PBS. Twenty cream-colored larvae for each group were selected and 5 µl of bacterial suspension was directly injected into the left proleg. One hour later, 5 µl of PBS containing 100 MOI of PICI-based EC-CapsidCas13a_*bla*_IMP-1_, EC-CapsidCas13a_null, or PBS only were injected into the same site where bacteria solution was injected. Thereafter, the larvae were transferred to a 37°C incubator, and the survival was observed up to 3 days. The larvae that did not respond to stimulation with needles or whose bodies deformed were counted as dead. Kaplan-Meier survival curves were then generated from data of three independent experiments and analyzed with log-rank test using software EZR (http://www.jichi.ac.jp/saitama-sct/SaitamaHP.files/statmedEN.html).

## Supporting information

Supplementary data combined

## Data availability

The data that support the findings in this study are available upon reasonable request from the corresponding authors.

## Acknowledgments

We thank Dr. Feng Zhang of the Massachusetts Institute of Technology, Dr. Luciano A Marraffini of The Rockefeller University, Dr. David Bikard of The Institut Pasteur and Dr. Timothy K. Lu of the Massachusetts Institute of Technology for their kindly gifting plasmids used in this study. We thank Hibiki Research Group for Clinical Microbiology for kindly providing Carbapenem-resistance *E. coli* strains R10-61 and R10-79. We thank Dr. Keiichi Hiramatsu of Juntendo University for kindly providing USA300 strain. Funding: This work was supported by the Japan Agency for Medical Research and Development J-PRIDE (grant No. JP17fm0208028, JP18fm0208028, and JP19fm0208028 to LC), JSPS KAKENHI (Grant No. 18K15149 to KK, 15H05654 and 19K08960 to SW, 17K15691 to YS, 19K15740 to MK and 17K19570 to LC), Mochida Memorial Foundation for Medical and Pharmaceutical Research (KK), and the Takeda Science Foundation (SW, LC). The funders had no role in the study design, data collection and analysis, decision to publish, or preparation of the manuscript.

## Contributions

KK, LC designed the study, analyzed the data, and wrote the manuscript. XT, RIC contributed to acquisition, analysis and interpretation of data, and assisted the preparation of the manuscript. JRP contributed to design of study, interpretation of data and assisted the preparation of the manuscript. All other authors contributed to data collection and interpretation, and critically revised the manuscript. All authors approved the final version of the manuscript and agreed for all aspects of the work in ensuring that questions related to the accuracy or integrity of any part of the work are appropriately investigated and resolved.

