## Supplementary data combined for "Development of CRSIPR-Cas13a-based antimicrobials capable of sequence-specific killing of target bacteria"

**Integrated supplementary informatnion:**

Supplementary Figures 1–8

Supplementary Tables 1–4

**Supplementary Figure 1**

Variation in bacterial killing activity of CRISPR-Cas13a system depending on the spacer sequence. The *E. coli* MC1061 derivatives harboring plasmid with expression of different AMR genes (pKLC26_*rfp*, pKLC26_*bla*$_{IMP-1}$, pKLC26_*bla*$_{OXA-48}$, pKLC26_*bla*$_{NDM-1}$, pKLC26_*bla*$_{KPC-2}$, pKLC26_*mcr-1*, and pKLC26_*mcr-2*) were transformed with CRISPR-Cas13a expression vector pKLC21(s) carrying different spacer sequences. The transformants were then plated on agar containing KM and CP, and incubated at 37° C for 12 hours, followed by counting colonies formed on the plates. **a**, The bar graph demonstrated that *E. coli* MC1061 derivatives, even though carrying the same target gene, showed distinctly different survival depending on the spacer sequence of the CRISPR-Cas13a system as shown in colony ratio. **b**, Representative plates for calculating colony ratio.

**Supplementary Figure 2**

Evaluation of spacer sequences targeting *bla*$_{IMP-1}$. **a**, pKLC21_*bla*$_{IMP-1}$ library with 121 different spacers targeting *bla*$_{IMP-1}$ were constructed and transformed into *E. coli* with and without *bla*$_{IMP-1}$. Plasmids retained in surviving cells were extracted and sequenced to identify the most effective spacer sequence. **b**, The sequence reads of spacers counted were normalized with that of non-targeting spacers (spacer_null) to calculate relative read number. Blue bar represents relative read number of the plasmids extracted from *E. coli* without carrying *bla*$_{IMP-1}$, while red bar represents the relative read number of the plasmids extracted from *E. coli* carrying *bla*$_{IMP-1}$. **c**, Depletion efficiency of each spacer was calculated by dividing the number of reads of blue bar by that of red bar, and the 10 spacer sequences with the highest depletion efficiency were listed.

**Supplementary Figure 3**

Schematic illustration of the construction of PICI-based EC-CapsidCas13a(s) in *E. coli* (see Methods).

**Supplementary Figure 4**

M13 phage-based EC-CapsidCas13a as a detection tool for specific bacterial gene. **a**, Schematic model illustrated bacterial gene detection method. **b**, In this assay, bacteria mixed with soft agar were poured onto 2 types of bottom agar plates, the LB plates and the LB-KM plates, followed by spotting of serially diluted EC-CapsidCas13a(s) carrying KM resistance gene (EC-CapsidCas13a::KanR(s)) onto the surface of the plates. The killing of *E. coli* NEB 5-alpha F'I$^{q}$ expressing *bla*$_{IMP-1}$ on LB plate **b** and LB-KM plate **c** was clearly seen when the EC-CapsidCas13a::KanR_*bla*$_{IMP-1}$ targeting *bla*$_{IMP-1}$ were spotted, indicating that M13-based EC-

CapsidCas13a could detect the gene of interest. The detection ability was enhanced by at least >1000 times by inserting the *kanR* gene into the EC-CapsidCas13a_$bla_{_IMP-1}$ *to* generate EC-CapsidCas13a::KanR_$bla_{IMP-1}$ **c**, compared with the results of directly using EC-CapsidCas13a_$bla_{IMP-1}$ **b**. **d**, The applicability of M13-based EC-CapsidCas13a::KanR(s) for detection of different kind of carbapenem resistance genes ($bla_{IMP-1}$, $bla_{OXA-48}$, and $bla_{VIM-2}$) located on both plasmid **d** and chromosome **e** was tested. Note that resistance genes on both plasmid and chromosome can be clearly detected and identified.

**Supplementary Figure 5**

Comparison of two kinds of bacterial lawns used for bacterial gene detection. Schematic model illustrated bacterial gene detection using plate prepared by soft agar overlay method **a**, and the detection results of genes on plasmid **b** and chromosome **c** were presented. The detection sensitivity was slightly lower, when the bacterial lawns were prepared by swabbing bacteria directly onto the agar plate **d** in detection of bacterial genes both on plasmid **e** and chromosome **f**, but not significantly different.

**Supplementary Figure 6**

Activity of CRISPR-Cas13a system in *Staphylococcus aureus.* **a**, A laboratory strain of *S. aureus* RN4220 carrying *rpsE* but not *ermC* and *mecA* was transformed with a series of pKLC4(s), which carry CRISPR-Cas13a targeting *rpsE* (20 points)*, ermC* (one point) and *mecA* (one point). **b**, The transformants were cultured on the TSA plate containing CP and CFU of survived cells were counted to test the sequence-specific killing activity of CRISPR-Cas13a against *S. aureus* by recognizing corresponding genes. To properly design CRISPR-Cas13a_*mecA* targeting *mecA* of MRSA, at first, pKLC4_*mecA*(s) with 20 different spacers against *mecA* were constructed and transformed into DH5$\alpha$ carrying anhydrotetracycline-inducible *mecA* expression vector to establish a spacer evaluation system **c**. Note that sequence-specific bacterial killing activity of CRISPR-Cas13a programed to target *mecA* was clearly dependent on the level of target gene expression **d**. Then, the 20 spacers designed against *mecA* were tested for their efficiency on mediating bacterial killing **e**.

**Supplementary Figure 7**

Schematic illustration of the SaPI-based SA-CapsidCas13a construction.

**Supplementary Figure 8**

Sequence-specific bacterial killing activity of SA-CapsidCas13a_*mecA* targeting *mecA* of methicillin-resistant *Staphylococcus aureus* (MRSA). Top agar containing bacterial lawn of MRSA USA300 and SaPI-based SA-CapsidCas13a::TetR_*mecA* were poured on the surface of TSA plates containing Tet. The plates were incubated until Tet-resistant colonies appear.

### Supplementary Fig. 1

**a**

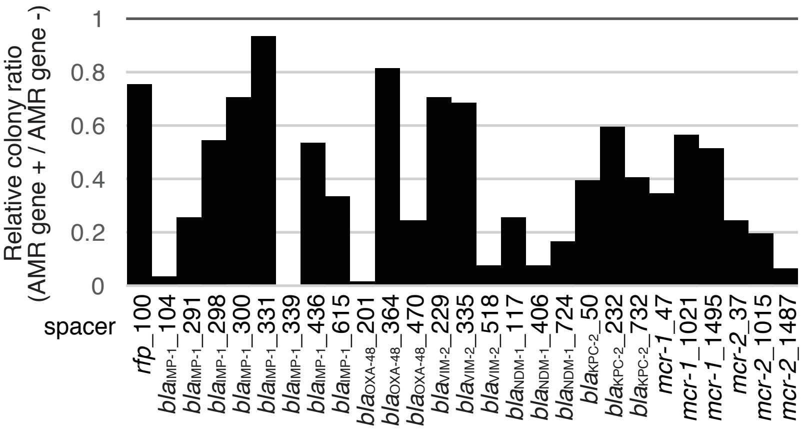


**b**

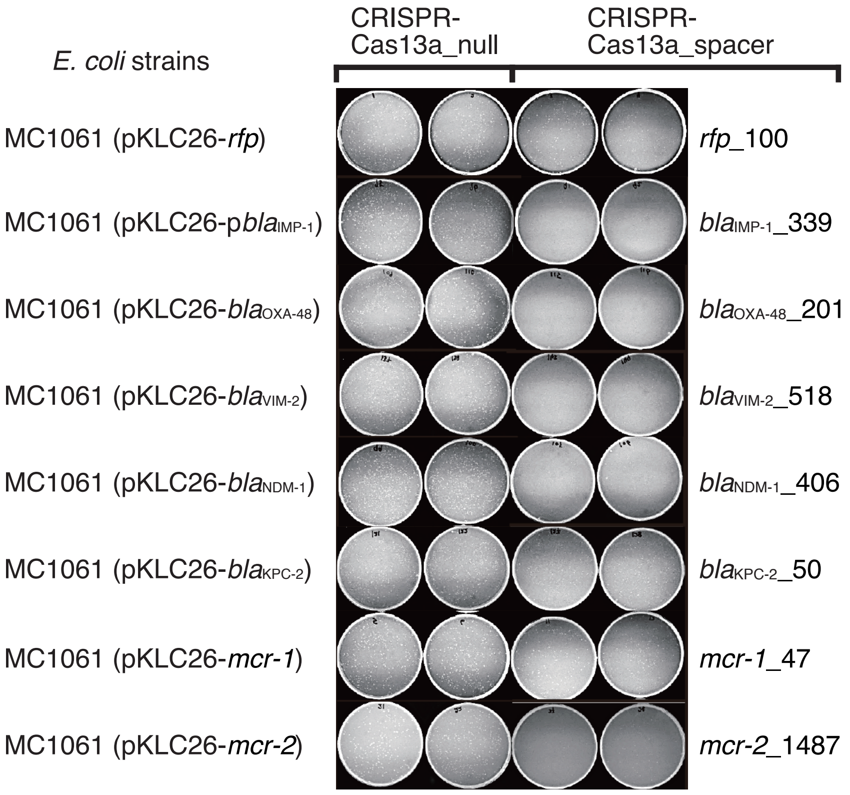

### Supplementary Fig. 2

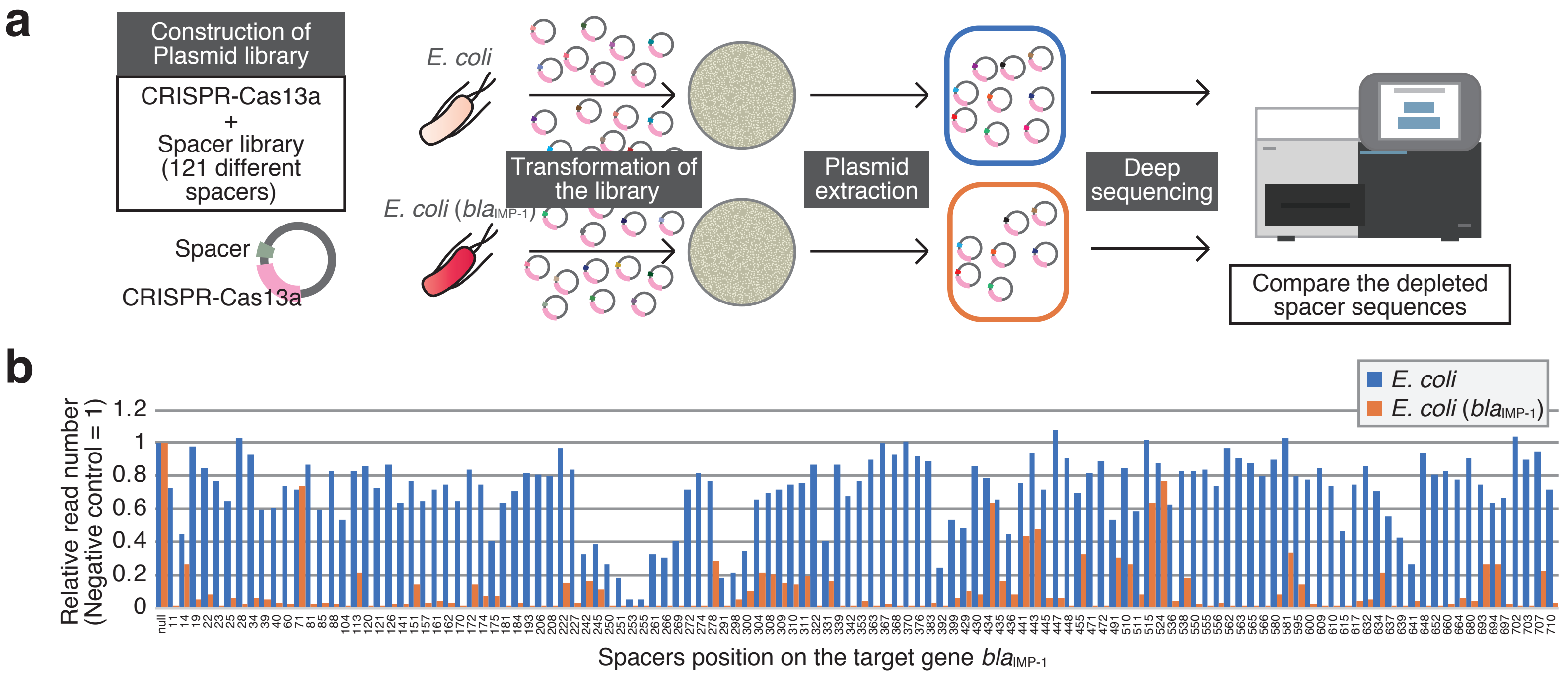


c

| Position in *bla*IMP-1 | Spacer sequence | Depletion efficiency |
|---|---|---|
| IMP-1_563 | GACTTTGGCCAAGCTTCTATATTTGCGT | 364.1 |
| IMP-1_566 | GCGGACTTTGGCCAAGCTTCTATATTTG | 256.8 |
| IMP-1_702 | TTTTGATGGTTTTTTACTTTCGTTTAAC | 236.9 |
| IMP-1_562 | ACTTTGGCCAAGCTTCTATATTTGCGTC | 220.8 |
| IMP-1_370 | TGGCTTGAACCTTACCGTCTTTTTTAAG | 201.4 |
| IMP-1_565 | CGGACTTTGGCCAAGCTTCTATATTTGC | 175.4 |
| IMP-1_193 | TATCTTTAGCCGTAAATGGAGTGTCAAT | 144.6 |
| IMP-1_274 | TGCTGTCGCTATGAAAATGAGAGGAAAT | 144.6 |
| IMP-1_648 | TTTCAAGAGTGATGCGTCTCCAACTTCA | 136.2 |
| IMP-1_11 | CAAAACAAAAATATAAAGAATACAGATA | 128.8 |
| IMP-1_455 | TTATCTGGAGTGTGTCCCGGGCCTGGAT | 2.2 |
| IMP-1_443 | TGTCCCGGGCCTGGATAAAAAACTTCAA | 2.0 |
| IMP-1_242 | CCTTTTATTTTATAGCCACGCTCCACAA | 2.0 |
| IMP-1_491 | AATAATATTTTCCTTTCAGGCAACCAAA | 1.8 |
| IMP-1_441 | TCCCGGGCCTGGATAAAAAACTTCAATT | 1.7 |
| IMP-1_14 | CTGCAAAACAAAAATATAAAGAATACAG | 1.7 |
| IMP-1_515 | TACGGTTTAATAAAACAACCACCGAATA | 1.6 |
| IMP-1_434 | CCTGGATAAAAAACTTCAATTTTATTTT | 1.2 |
| IMP-1_524 | CCTAAACCGTACGGTTTAATAAAACAAC | 1.1 |
| IMP-1_71 | CCTTCATCAAGCTTTTCAATTTTTAAAT | 1.0 |

### Supplementary Fig. 3

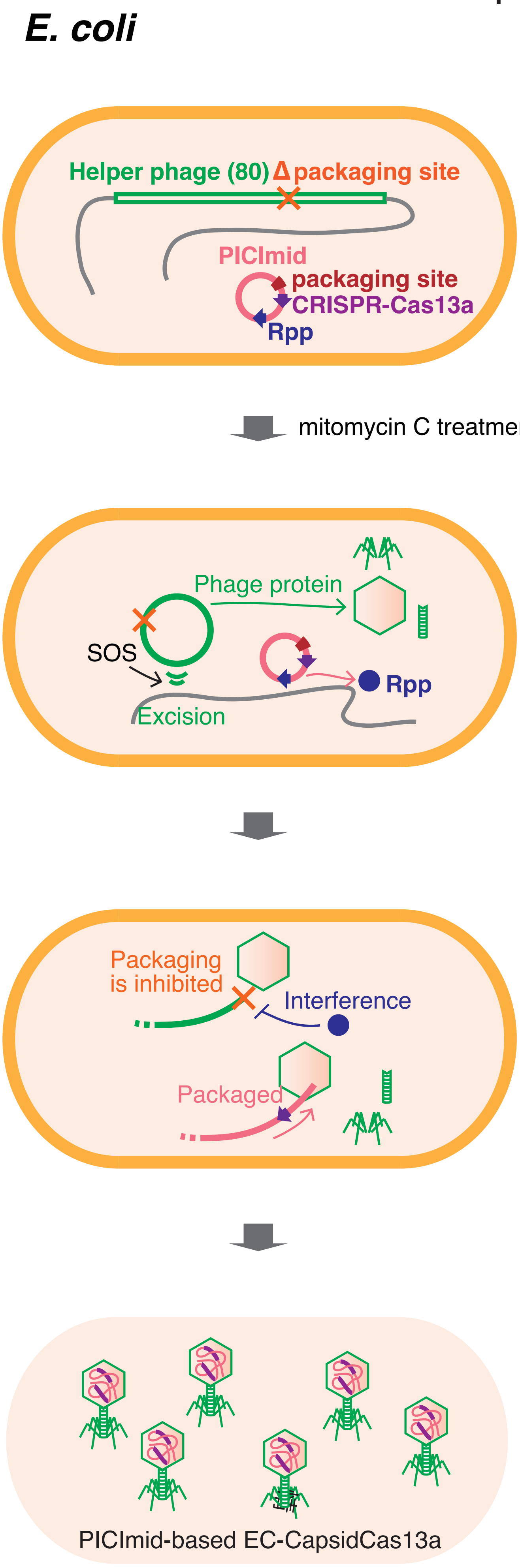

### Supplementary Fig. 4

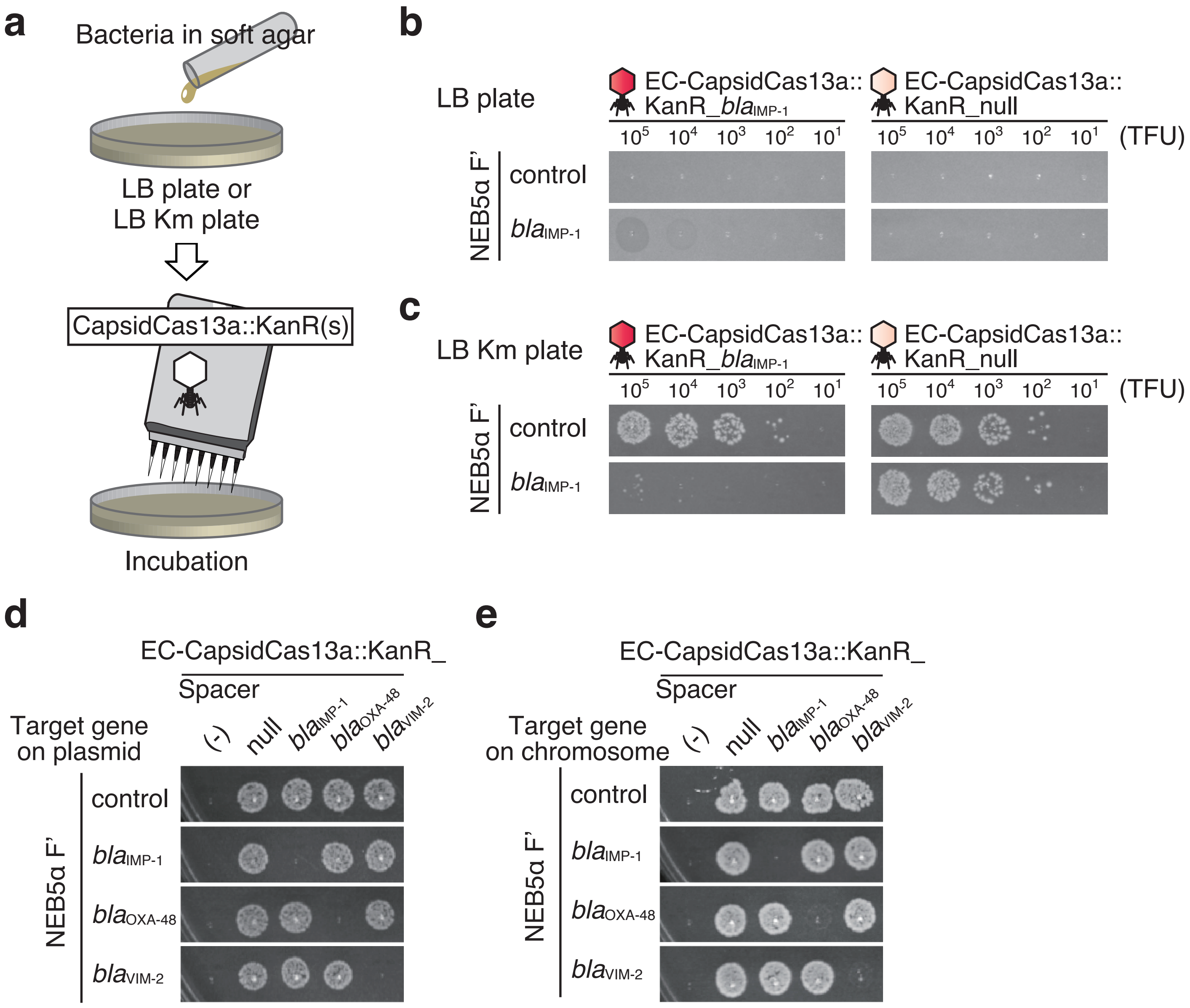

### Supplementary Fig. 5

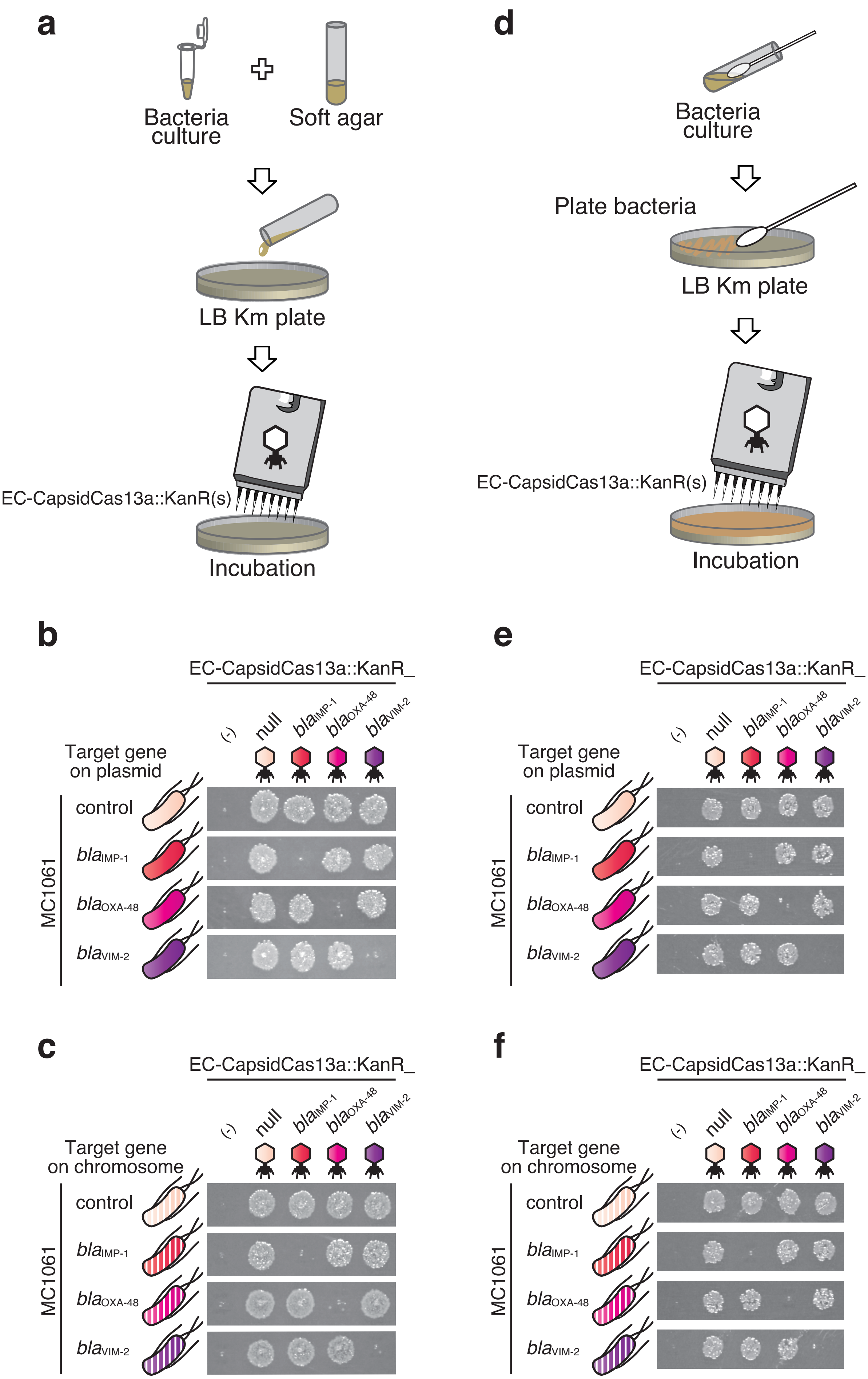

### Supplementary Fig. 6

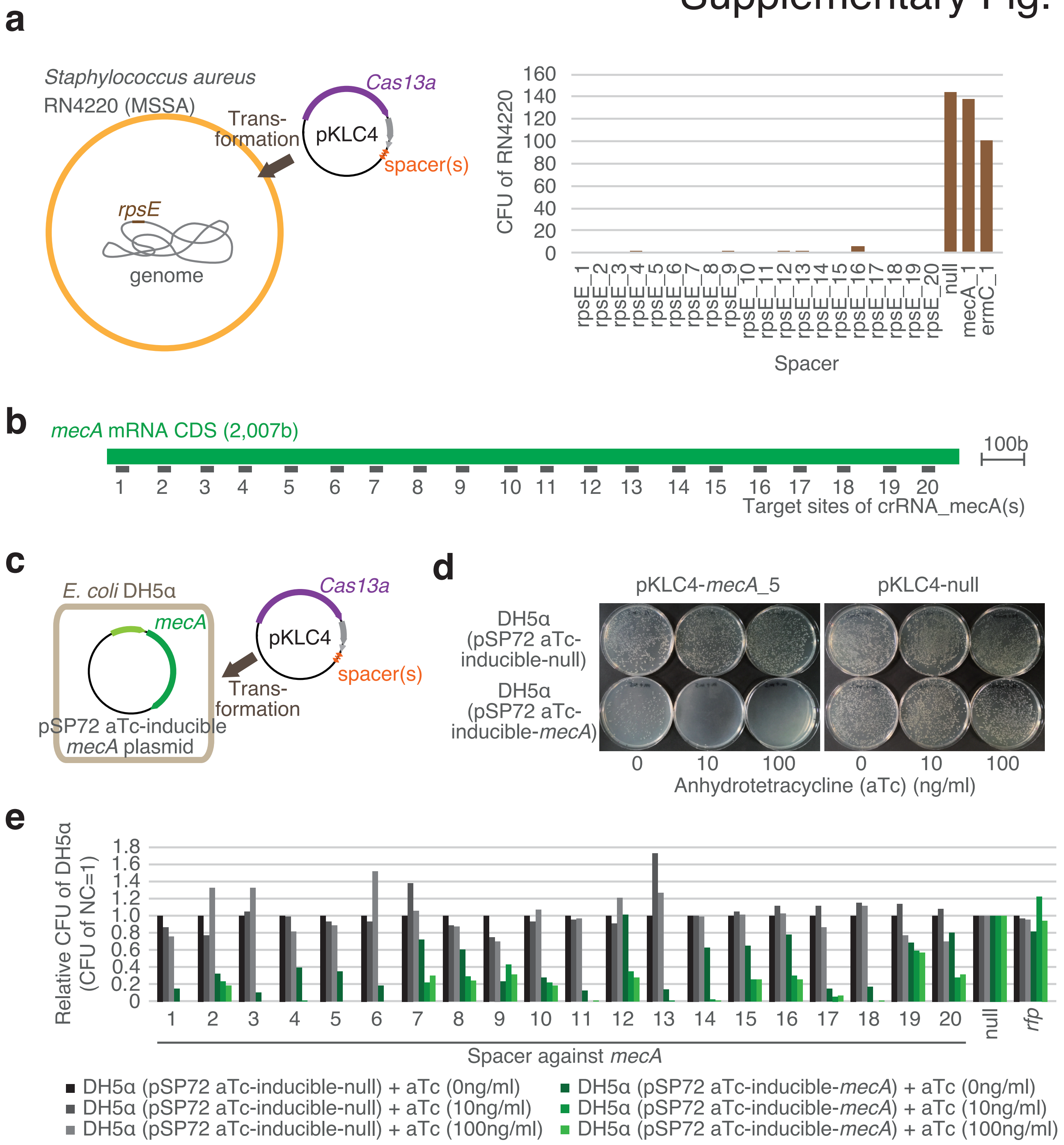

### Supplementary Fig. 7

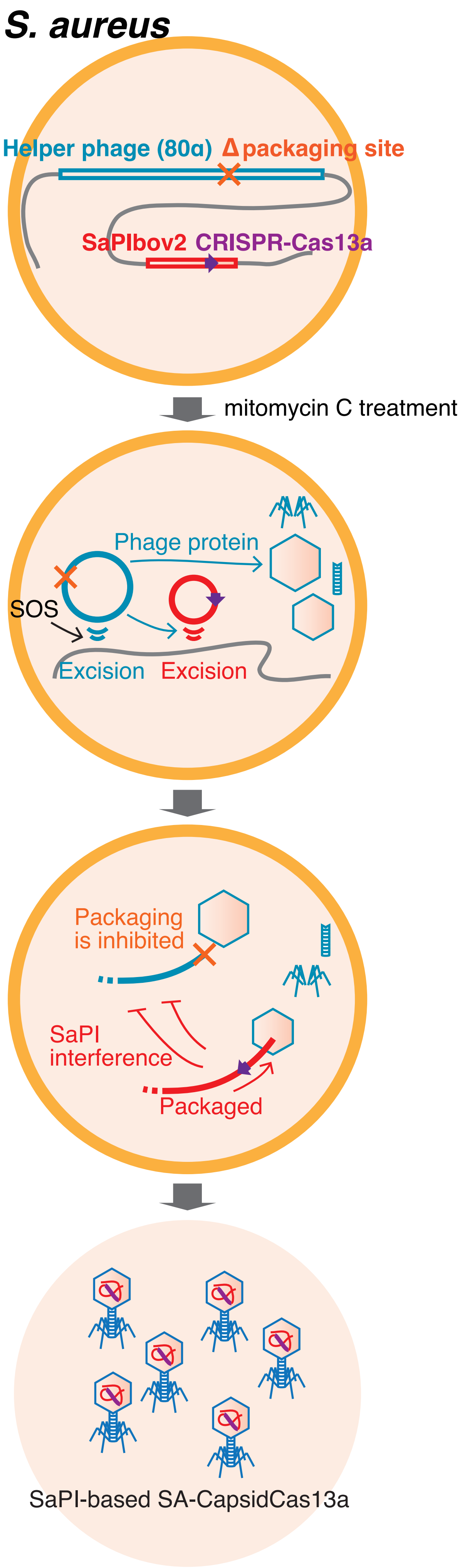

### Supplementary Fig. 8

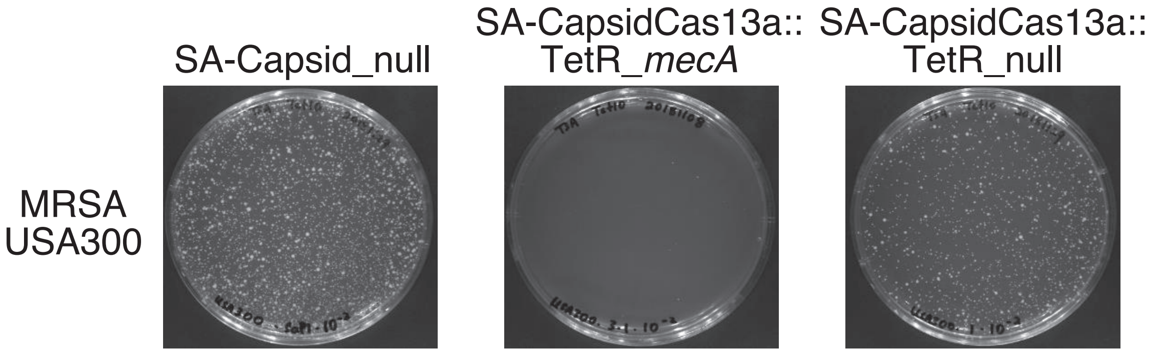

**Table S1. List of Bacterial strains used in this study**

| Bacterial strain | Description | Origin |
|---|---|---|
| MC1061 | hsdR, mcrB, araD139, Δ(araABC-leu)7679, ΔlacX74, galU, galK, rpsL, thi | Casadaban et al., 1980[a] |
| NEB5-alpha F' $I^q$ | F' proA+B+ lacIq Δ(lacZ)M15 zzf::Tn10 (TetR) / fhuA2Δ(argF-lacZ)U169 phoA glnV44 Φ80Δ(lacZ)M15 gyrA96 recA1 relA1 endA1 thi-1 hsdR17 | NEB biolabs[b] |
| NEB5α F' (pPBAD-control) | NEB5-alpha F' lq transformed with pKLC23 | in this study |
| NEB5α F' (pPBAD-*bla*$_{\text{IMP-1}}$) | NEB5-alpha F' lq transformed with pKLC23-*bla*$_{\text{IMP-1}}$ | in this study |
| NEB5α F' (pControl) | NEB5-alpha F' lq transformed with pKLC26 | in this study |
| NEB5α F' (p*bla*$_{\text{IMP-1}}$) | NEB5-alpha F' lq transformed with pKLC26-*bla*$_{\text{IMP-1}}$ | in this study |
| NEB5α F' (p*bla*$_{\text{OXA-48}}$) | NEB5-alpha F' lq transformed with pKLC26-*bla*$_{\text{OXA-48}}$ | in this study |
| NEB5α F' (p*bla*$_{\text{VIM-2}}$) | NEB5-alpha F' lq transformed with pKLC26-*bla*$_{\text{VIM-2}}$ | in this study |
| NEB5α F' (p*bla*$_{\text{NDM-1}}$) | NEB5-alpha F' lq transformed with pKLC26-*bla*$_{\text{NDM-1}}$ | in this study |
| NEB5α F' (p*bla*$_{\text{KPC-2}}$) | NEB5-alpha F' lq transformed with pKLC26-*bla*$_{\text{KPC-2}}$ | in this study |
| NEB5α F' (p*mcr-1*) | NEB5-alpha F' lq transformed with pKLC26-*mcr-1* | in this study |
| NEB5α F' (p*mcr-2*) | NEB5-alpha F' lq transformed with pKLC26-*mcr-2* | in this study |
| NEB5α F' *bla*$_{\text{NDM-1}}$ in chromosome | MC1061 *bla*$_{\text{NDM-1}}$ sequence was inserted in the chromosome | in this study |
| R10-61 | Carbapenem-resistant E. coli | Hibiki Research Group for Clinical Microbiology |
| R10-79 | Carbapenem-resistant E. coli | Hibiki Research Group for Clinical Microbiology |
| MC1061 (pControl) | MC1061 transformed with pKLC26 | in this study |
| MC1061 (p*bla*$_{\text{IMP-1}}$) | MC1061 transformed with pKLC26-*bla*$_{\text{IMP-1}}$ | in this study |
| MC1061 (p*bla*$_{\text{OXA-48}}$) | MC1061 transformed with pKLC26-*bla*$_{\text{OXA-48}}$ | in this study |
| MC1061 (p*bla*$_{\text{VIM-2}}$) | MC1061 transformed with pKLC26-*bla*$_{\text{VIM-2}}$ | in this study |
| MC1061 *bla*$_{\text{IMP-1}}$ in chromosome | MC1061 *bla*$_{\text{IMP-1}}$ sequence was inserted in the chromosome | in this study |
| MC1061 *bla*$_{\text{OXA-48}}$ in chromosome | MC1061 *bla*$_{\text{OXA-48}}$ sequence was inserted in the chromosome | in this study |
| MC1061 *bla*$_{\text{VIM-2}}$ in chromosome | MC1061 *bla*$_{\text{VIM-2}}$ sequence was inserted in the chromosome | in this study |
| MC1061 (p*stx-1*) | MC1061 transformed with pKLC26-*stx-1* (partial) | in this study |
| MC1061 (p*stx-2*) | MC1061 transformed with pKLC26-*stx-2* (partial) | in this study |
| RN4220 | Laboratory strain of *S. aureus* | Kreiswirth BN et al., 1983[c] |
| USA300 | Methicillin-resistant *S. aureus* (MRSA), FPR3757 | Diep BA et al., 2006[d] |
| USA300 Δ*mecA* | *mecA* deletion mutant of USA300 | in this study |

a. M. J. Casadaban, S. N. Cohen. Analysis of gene control signals by DNA fusion and cloning in Escherichia coli. J Mol Biol. (1980) doi:10.1016/0022-2836(80)90283-1, Lucigen (https://www.lucigen.com/)

b. Neu England Biolabs (https://international.neb.com/)

c. B. N. Kreiswirth, S. Löfdahl, M. J. Betley, M. O'Reilly, P. M. Schlievert, M. S. Bergdoll, R. P. Novick, The toxic shock syndrome exotoxin structural gene is not detectably transmitted by a prophage. Nature (1983), doi:10.1038/305709a0

d. B.A. Diep, S.R. Gill, R. F. Chang, T. H. Phan, J. H. Chen, M. G. Davidson, F. Lin, J. Lin, H. A. Carleton, E. F. Mongodin, G. F. Sensabaugh, F. Perdreau-Remington, Complete genome sequence of USA300, an epidemic clone of community-acquired meticillin-resistant Staphylococcus aureus. Lancet (2006) doi:10.1016/S0140-6736(06)68231-7

**Table S2. List of Vectors used in this study**

| Vectetor | Purpose | Discription | Origin |
|---|---|---|---|
| pSP72 | Construction of plasmid | Cloning vector, multiple cloning site, β-lactamase, SP6 promoter, T7 promoter | Promega[a] |
| pC003 | | Cas13 expression vector, Cas1, Cas2, p15A ori, TcR, CAT | From Dr. Feng Zhang[b] |
| pDB114 | | Cas9 expression vector in S. aureus, CAT, rep from pC194 | From Dr. Luciano A. Marraffini[c] |
| pRC319 | | Cas9 expression vector, pBR322 ori, F1 ori, KanR, crRNA for blaNDM-1 | From Dr. Timothy Lu[d] |
| pKLC3.0 | | Shuttle vector for Cas13a expression in E. coli and S. aureus, p15A ori for E. coli, KanR (E. coli), CAT (S. aureus), rep for S. aureus (pC194) | This study |
| pKLC4 | | Shuttle vector for CRISPR-Cas13a expression in E. coli and S. aureus, p15A ori for E. coli, KanR (E. coli), CAT (S. aureus), rep for S. aureus (pC194) in this study | This study |
| pSP72 aTc-inducible | Tetracycline-inducible target gene expression vector | Expression vector under the control of tetracycline, pBR322 ori, beta-lactamase | This study |
| pSP72 aTc-inducible *mecA* | | *mecA* expression vector under the control of tetracycline, pBR322 ori, beta-lactamase | This study |
| pSP72 aTc-inducible *bla* IMP-1 | | *bla* IMP-1 expression vector under the control of tetracycline, pBR322 ori, beta-lactamase | This study |
| pKLC21 | Construction of M13-based CapsidCas13a::KanR | CRISPR-Cas13a expression vector in E. coli, ColE1 ori, KanR, f1 ori | This study |
| pKLC21_*blaI* MP-1_339 | | CRISPR-Cas13a (target: *bla* IMP-1_339) expression vector in E. coli, ColE1 ori, KanR, f1 ori | This study |
| pKLC21_*blaI* MP-1_563 | | CRISPR-Cas13a (target: *bla* IMP-1_563) expression vector in E. coli, ColE1 ori, KanR, f1 ori | This study |
| pKLC21_*bla* oxa-48_201 | | CRISPR-Cas13a (target: *bla* OXA-48_201) expression vector in E. coli, ColE1 ori, KanR, f1 ori | This study |
| pKLC21_*bla* VIM-2_518 | | CRISPR-Cas13a (target: *bla* VIM-2_518) expression vector in E. coli, ColE1 ori, KanR, f1 ori | This study |
| pKLC21_*bla* NDM-1_406 | | CRISPR-Cas13a (target: *bla* NDM-1_406) expression vector in E. coli, ColE1 ori, KanR, f1 ori | This study |
| pKLC21_*bla* KPC-2_50 | | CRISPR-Cas13a (target: *bla* KPC-2_50) expression vector in E. coli, ColE1 ori, KanR, f1 ori | This study |
| pKLC21_*mcr-1* _47 | | CRISPR-Cas13a (target: *mcr-1* _50) expression vector in E. coli, ColE1 ori, KanR, f1 ori | This study |
| pKLC21_*mcr-2* _1487 | | CRISPR-Cas13a (target: *mcr-2* _1487) expression vector in E. coli, ColE1 ori, KanR, f1 ori | This study |
| pKLC23 | Arabinose-inducible target gene expression vector | Antibiotics resistance gene expression vector, p15A ori, PBAD ptomoter, CAT | This study |
| pKLC25 | Construction of M13 phage | M13 helper phage vector, p15A ori, CAT, f1 ori is deleted, E. coli transformed with this vector grow slowly | This study |
| pKLC26 | Constitutive target gene expression vector | Target gene expression vector, p15A ori, intI1 promoter (native promoter of *bla* IMP-1), CAT | This study |
| pKLC26_*blaI* MP-1 | | *bla* IMP-1 expression vector, p15A ori, intI1 promoter (native promoter of *bla* IMP-1), CAT | This study |
| pKLC26_*bla* oxa-48 | | *bla* OXA-48 expression vector, p15A ori, intI1 promoter (native promoter of *bla* IMP-1), CAT | This study |
| pKLC26_*bla* VIM-2 | | *bla* VIM-2 expression vector, p15A ori, intI1 promoter (native promoter of *bla* IMP-1), CAT | This study |
| pKLC26_*bla* NDM-1 | | *bla* NDM-1 expression vector, p15A ori, intI1 promoter (native promoter of *bla* IMP-1), CAT | This study |
| pKLC26_*bla* KPC-2 | | *bla* KPC-2 expression vector, p15A ori, intI1 promoter (native promoter of *bla* IMP-1), CAT | This study |
| pKLC26_*mcr-1* | | *mcr-1* expression vector, p15A ori, intI1 promoter (native promoter of *bla* IMP-1), CAT | This study |
| pKLC26_*mcr-2* | | *mcr-2* expression vector, p15A ori, intI1 promoter (native promoter of *bla* IMP-1), CAT | This study |
| pKLC26_*mcr-3* | | *mcr-3* expression vector, p15A ori, intI1 promoter (native promoter of *bla* IMP-1), CAT | This study |
| pKLC26_*mcr-4* | | *mcr-4* expression vector, p15A ori, intI1 promoter (native promoter of *bla* IMP-1), CAT | This study |
| pKLC26_*mcr-5* | | *mcr-5* expression vector, p15A ori, intI1 promoter (native promoter of *bla* IMP-1), CAT | This study |
| pKLC26_*stx-1* (partial) | | *stx-1* (partial) expression vector, p15A ori, intI1 promoter (native promoter of *bla* IMP-1), CAT | This study |
| pKLC26_*stx-2* (partial) | | *stx-2(partial)* expression vector, p15A ori, intI1 promoter (native promoter of *bla* IMP-1), CAT | This study |

| | | | |
|---|---|---|---|
| pKLC31 | Construction of PICI-based CapsidCas13a::KanR | PICI-construction vector, CRISPR-Cas13a, KanR, pBR322 ori, aTc-inducible c1501/c1502/c1503 | This study |
| pKLC31_*blaI* MP-1_339 | | PICI-construction vector, CRISPR-Cas13a (target: *bla* IMP-1_339), KanR, pBR322 ori, aTc-inducible c1501/c1502/c1503 | This study |
| pKLC31_*blaI* MP-1_563 | | PICI-construction vector, CRISPR-Cas13a (target: *bla* IMP-1_563), KanR, pBR322 ori, aTc-inducible c1501/c1502/c1503 | This study |
| pKLC31_*bla* oxa-48_201 | | PICI-construction vector, CRISPR-Cas13a (target: *bla* OXA-48_201), KanR, pBR322 ori, aTc-inducible c1501/c1502/c1503 | This study |
| pKLC31_*bla* VIM-2_518 | | PICI-construction vector, CRISPR-Cas13a (target: *bla* VIM-2_518), KanR, pBR322 ori, aTc-inducible c1501/c1502/c1503 | This study |
| pKLC31_*bla* NDM-1_406 | | PICI-construction vector, CRISPR-Cas13a (target: *bla* NDM-1_406), KanR, pBR322 ori, aTc-inducible c1501/c1502/c1503 | This study |
| pKLC31_*bla* KPC-2_50 | | PICI-construction vector, CRISPR-Cas13a (target: *bla* KPC-2_50), KanR, pBR322 ori, aTc-inducible c1501/c1502/c1503 | This study |
| pKLC31_*mcr-1* _47 | | PICI-construction vector, CRISPR-Cas13a (target: *mcr-1* _47), KanR, pBR322 ori, aTc-inducible c1501/c1502/c1503 | This study |
| pKLC31_*mcr-2* _1487 | | PICI-construction vector, CRISPR-Cas13a (target: *mcr-2* _1487), KanR, pBR322 ori, aTc-inducible c1501/c1502/c1503 | This study |
| pKLC31_stx-*1* _640 | | PICI-construction vector, CRISPR-Cas13a (target: *mcr-1* _47), KanR, pBR322 ori, aTc-inducible c1501/c1502/c1503 | This study |
| pKLC31_stx-*2* _640 | | PICI-construction vector, CRISPR-Cas13a (target: *mcr-2* _1487), KanR, pBR322 ori, aTc-inducible c1501/c1502/c1503 | This study |
| pKLC44 | Construction of PICI-based CapsidCas13a::HygroR | PICI-construction vector, CRISPR-Cas13a, HygroR, pBR322 ori, aTc-inducible c1501/c1502/c1503 | This study |
| pKLC44_*blaI* MP-1_563 | | PICI-construction vector, CRISPR-Cas13a (target: *bla* IMP-1_563), HygroR, pBR322 ori, aTc-inducible c1501/c1502/c1503 | This study |
| pKLC44_*bla* NDM-1_406 | | PICI-construction vector, CRISPR-Cas13a (target: *bla* NDM-1_406), HygroR, pBR322 ori, aTc-inducible c1501/c1502/c1503 | This study |
| pC003-*mecA* 5 | Construction of *mecA* -targeting Cas13a | pC003 loaded with optimized crRNA targeting *mecA* | This study |
| pC003-*mecA* 5-ΔCas1/2 | | pC003-*mecA* 5 with deleted Cas1 and Cas2 | This study |
| pIMAY | Construction of SaPI-based CapsidCas13a::TetR | *E. coli* /staphylococcal temperature-sensitive plasmid, ori for *E. coli* p15A, P*help-cat* , anti-*secY* , temperature-sensitive replicon for Gram-positive bacteria (*repBCAD* ) | Monk *et al.* 2012[e] |
| pIMAY-BAPup/down | | pIMAY carrying 5' and 3' flanking regions of *bap* gene | This study |
| pIMAY-LshCas13a-SP*mecA* | | pIMAY-BAPup/down with LshCas13a and crRNA targeting *mecA* inserted in between 5' and 3' flanking regions of *bap* gene | This study |
| pIMAY-LshCas13a-null | | pIMAY-BAPup/down with only LshCas13a inserted in between 5' ar | This study |
| pIMAY-KO*mecA* | | pIMAY carrying 5' and 3' flanking regions of *mecA* gene | This study |

a. Thermo Fisher Scientific (Promega) (https://www.fishersci.ca/shop/products/promega-psp72-vector/prp2191)

b. O. O. Abudayyeh, J. S. Gootenberg, S. Konermann, J. Joung, I. M. Slaymaker, D. B. T. Cox, S. Shmakov, K. S. Makarova, E. Semenova, L. Minakhin, K. Severinov, A. Regev, E. S. Lander, E. v. Koonin, F. Zhang, C2c2 is a single-component programmable RNA-guided RNA-targeting CRISPR effector. Science (2016), doi:10.1126/science.aaf5573.

c. D. Bikard, C. W. Euler, W. Jiang, P. M. Nussenzweig, G. W. Goldberg, X. Duportet, V. A. Fischetti, L. A. Marraffini, Exploiting CRISPR-cas nucleases to produce sequence-specific antimicrobials. Nature Biotechnology (2014), doi:10.1038/nbt.3043.

d. R. J. Citorik, M. Mimee, T. K. Lu, Sequence-specific antimicrobials using efficiently delivered RNA-guided nucleases. Nature Biotechnology (2014), doi:10.1038/nbt.3011.

e. I. R. Monk, I. M. Shah, M. Xu, M. W. Tan, T. J. Foster, Transforming the untransformable: application of direct transformation to manipulate genetically Staphylococcus aureus and Staphylococcus epidermidis. MBio (2012), doi: 10.1128/mBio.00277-11.

**Table S3. List of Primers used in this study**

| Purpose | Primer name | Sequence |
|---|---|---|
| Deletion of Cas1/2 from pC003 | Cas1Cas2 del SacI-f | ATATGAGCTCATGGGAGAAAAAATTTCACAAAAC |
| | Cas1Cas2 del SacI-r | ATATGAGCTCTCATTCTTATAACGTATCATTCG |
| Construction of pKLC3.0 | InF3.0 KanR-f | gcattaaagctcgtttaacagcGTTTTTAGTTGAAAGCTAACTTC |
| | InF3.0 KanR-r | ttccacattttcccaCAGTGAATTGGAGTTCGTC |
| | InF3.0 SArep_CAT-f | acactccgctagcgcCAAACGAAAATTGGATAAAGTG |
| | InF3.0 SArep_CAT-r | gctgttaaacgagctttaatgcCGTTTGTTGAACTAATGGGTG |
| | InF3.0 p15A ori_Cas13-f | TGGGAAAATGTGGAATTTGAAAC |
| | InF3.0 p15A ori_Cas13-r | GCGCTAGCGGAGTGTATACTG |
| Construction of pKLC21 | InF13 SalI PCR-f | tcttcaccctgtcgatgggaaaatgtggaatttg |
| | InF13 SmaI PCR-r | caggatcttctgcccaattaggctctagttagcct |
| | InF13 pRC319-f | gggcagaagatcctgcagg |
| | InF13 pRC319-r | tcgacagggtgaagacgaaag |
| Construction of pKLC21 IMP-1_104 | pC003 blaIMP-1_104-s | tatccATGTTCATACTTCGTTTGAAGAAGTTAA |
| | pC003 blaIMP-1_104-as | aaacTTAACTTCTTCAAACGAAGTATGAACATg |
| Construction of pKLC25 | M13KO7 PCR InFusion-f | cctattggttaaaaaatgagctg |
| | M13KO7 PCR InFusion-r | actatggttgctttgacgag |
| | pBAD33 PCR InFusion-f | caaagcaaccatagtgtagcaccaggcgtttaagg |
| | pBAD33 PCR InFusion-r | tttttaaccaataggcatcaccgatggggaagatc |
| Construction of pKLC23 | pBAD33 PCR InFusion-f | tcaCTCGAGcgaatttgctttcgaatttc |
| | pBAD33 PCR InFusion-r | tcaCTCGAGcaaaagagtttgtagaaacgc |
| Construction of pKLC26 | pKLC23 PCR InFusion-f | GAATTCgatcctctagagtc |
| | pKLC23 PCR InFusion-r | tgcttcgtccatttgacag |
| | Intl1pro PCR InFusion-f | caaatggacgaagcaTGACGCACACCGTGGAAAC |
| | Intl1pro PCR InFusion-r | tagaggatcGAATTCGAGAATGGATTTTGTGATGC |
| Construction of antibiotics resistant gene expression vector | InF18 pKLC26-f | GAATTCgatcctctagagtc |
| | InF18 pKLC26-r | GAGAATGGATTTTGTGATGC |
| | InF18 NDM-1-f | ACAAAATCCATTCTCATGGAATTGCCCAATATTATG |
| | InF18 NDM-1-r | tagaggatcGAATTCTCAGCGCAGCTTGTCGGC |
| | InF18 KPC-2-f | ACAAAATCCATTCTCATGTCACTGTATCGCCGTC |
| | InF18 KPC-2-r | tagaggatcGAATTCTTACTGCCCGTTGACGCC |
| | InF18 VIM-2-f | ACAAAATCCATTCTCATGTTCAAACTTTTGAGTAAG |
| | InF18 VIM-2-r | tagaggatcGAATTCCTACTCAACGACTGAGCG |
| | InF18 IMP-1-f | ACAAAATCCATTCTCATGAGCAAGTTATCTGTATTC |
| | InF18 IMP-1-r | tagaggatcGAATTCTTAGTTGCTTGGTTTTGATG |
| | InF18 OXA-48-f | ACAAAATCCATTCTCATGCGTGTATTAGCCTTATC |
| | InF18 OXA-48-r | tagaggatcGAATTCCTAGGGAATAATTTTTTCCTG |
| | InF18 mcr-1-f | ACAAAATCCATTCTCATGATGCAGCATACTTCTG |
| | InF18 mcr-1-r | tagaggatcGAATTCTCAGCGGATGAATGCGGTG |
| | InF18 mcr-2-f | ACAAAATCCATTCTCATGACATCACATCACTCTTG |
| | InF18 mcr-2-r | tagaggatcGAATTCTTACTGGATAAATGCCGC |
| | InF18 RFP-f | ACAAAATCCATTCTCatggcgagtagcgaagac |
| | InF18 RFP-r | tagaggatcGAATTCttaagcaccggtggagtg |
| Oligo DNA sequences for crRNA in E. coli | crR-IMP-1_104-as | tatccATGTTCATACTTCGTTTGAAGAAGTTAA |
| | crR-IMP-1_104-s | aaacTTAACTTCTTCAAACGAAGTATGAACATg |
| | crR-IMP-1_291-as | tatccTAGCGACAGCACGGGCGGAATAGAGTGG |
| | crR-IMP-1_291-s | aaacCCACTCTATTCCGCCCGTGCTGTCGCTAg |
| | crR-IMP-1_298-as | tatccAGCACGGGCGGAATAGAGTGGCTTAATT |
| | crR-IMP-1_298-s | aaacAATTAAGCCACTCTATTCCGCCCGTGCTg |
| | crR-IMP-1_300-as | tatccCACGGGCGGAATAGAGTGGCTTAATTCT |
| | crR-IMP-1_300-s | aaacAGAATTAAGCCACTCTATTCCGCCCGTGg |
| | crR-IMP-1_331-as | tatccTCTATCCCCACGTATGCATCTGAATTAA |
| | crR-IMP-1_331-s | aaacTTAATTCAGATGCATACGTGGGGATAGAg |
| | crR-IMP-1_339-as | tatccCACGTATGCATCTGAATTAACAAATGAA |
| | crR-IMP-1_339-s | aaacTTCATTTGTTAATTCAGATGCATACGTGg |
| | crR-IMP-1_436-as | tatccAATAAAATTGAAGTTTTTATCCAGGCC |
| | crR-IMP-1_436-s | aaacGGCCTGGATAAAAAACTTCAATTTTATTg |
| | crR-IMP-1_615-as | tatccTGGTAAGGCAAAACTGGTTGTTCCAAGT |
| | crR-IMP-1_615-s | aaacACTTGGAACAACCAGTTTTGCCTTACCAg |
| | crR-KPC-2_232-as | tatccGCTGTGCTGGCTCGCAGCCAGCAGCAGG |
| | crR-KPC-2_232-s | aaacCCTGCTGCTGGCTGCGAGCCAGCACAGCg |
| | crR-KPC-2_50-as | tatccTGGCTGGCTTTTCTGCCACCGCGCTGAC |
| | crR-KPC-2_50-s | aaacGTCAGCGCGGTGGCAGAAAAGCCAGCCAg |
| | crR-KPC-2_732-as | tatccTGACTATGCCGTCGTCTGGCCCACTGGG |
| | crR-KPC-2_732-s | aaacCCCAGTGGGCCAGACGACGGCATAGTCAg |
| | crR-NDM-1_117-as | tatccGGAAACTGGCGACCAACGGTTTGGCGAT |
| | crR-NDM-1_117-s | aaacATCGCCAAACCGTTGGTCGCCAGTTTCCg |
| | crR-NDM-1_406-as | tatccGGGATTGCGACTTATGCCAATGCGTTGT |
| | crR-NDM-1_406-s | aaacACAACGCATTGGCATAAGTCGCAATCCCg |
| | crR-NDM-1_724-as | tatccAAGGCCAGCATGATCGTGATGAGCCATT |
| | crR-NDM-1_724-s | aaacAATGGCTCATCACGATCATGCTGGCCTTg |
| | crR-OXA-48_201-as | tatccACCCGCATCTACCTTTAAAATTCCCAAT |
| | crR-OXA-48_201-s | aaacATTGGGAATTTTAAAGGTAGATGCGGGTg |
| | crR-OXA-48_364-as | tatccGTTTATCAAGAATTTGCCCGCCAAATTG |

| | | |
|---|---|---|
| | crR-OXA-48_364-s | aaacCAATTTGGCGGGCAAATTCTTGATAAACg |
| | crR-OXA-48_470-as | tatccGGCTCGACGGTGGTATTCGAATTTCGGC |
| | crR-OXA-48_470-s | aaacGCCGAAATTCGAATACCACCGTCGAGCCg |
| | crR-VIM-2_229-as | tatccGGTGATGAGTTGCTTTTGATTGATACAG |
| | crR-VIM-2_229-s | aaacCTGTATCAATCAAAAGCAACTCATCACCg |
| | crR-VIM-2_335-as | tatccCCACGCACTTTCATGACGACCGCGTCGG |
| | crR-VIM-2_335-s | aaacCCGACGCGGTCGTCATGAAAGTGCGTGGg |
| | crR-VIM-2_518-as | tatccTCTATCCTGGTGCTGCGCATTCGACCGA |
| | crR-VIM-2_518-s | aaacTCGGTCGAATGCGCAGCACCAGGATAGAg |
| | crR-mcr-1_1021-as | tatccAAAGCGCAATTTGCCGATTATAAATCCG |
| | crR-mcr-1_1021-s | aaacCGGATTTATAATCGGCAAATTGCGCTTTg |
| | crR-mcr-1_1495-as | tatccGATAAGCAAACTGGCATCACGCCAATGG |
| | crR-mcr-1_1495-s | aaacCCATTGGCGTGATGCCAGTTTGCTTATCg |
| | crR-mcr-1_47-as | tatccTTGTTCTTGTGGCGAGTGTTGCCGTTTT |
| | crR-mcr-1_47-s | aaacAAAACGGCAACACTCGCCACAAGAACAAg |
| | crR-mcr-2_1015-as | tatccGCCACGCAGTATTTTGATTATAAATCAG |
| | crR-mcr-2_1015-s | aaacCTGATTTATAATCAAAATACTGCGTGGCg |
| | crR-mcr-2_1487-as | tatccCAAATAATACGACATTCAAGCCAACTGC |
| | crR-mcr-2_1487-s | aaacGCAGTTGGCTTGAATGTCGTATTATTTGg |
| | crR-mcr-2_37-as | tatccAATCCTTTTGTGCTGATGGGTTTGGTGG |
| | crR-mcr-2_37-s | aaacCCACCAAACCCATCAGCACAAAAGGATTg |
| | crR-stx1_640-as | tatccCACCGGAAGAAGTGGAACTCACACTGAA |
| | crR-stx1_640-s | aaacTTCAGTGTGAGTTCCACTTCTTCCGGTGg |
| | crR-stx2_640-as | tatccCGCCGGGAGACGTGGACCTCACTCTGAA |
| | crR-stx2_640-s | aaacTTCAGAGTGAGGTCCACGTCTCCCGGCGg |
| Oligo DNA sequences for crRNA in S. aureus | BsaI rpsE 40-as | tatccTAGAGAAGAAGAGACGAAAGAATTTGAAGAA |
| | BsaI rpsE 40-s | aaacTTCTTCAAATTCTTTCGTCTCTTCTTCTCTAg |
| | BsaI rpsE 65-as | tatccGAAGAACGCGTTGTTACAATCAACCGTGTAG |
| | BsaI rpsE 65-s | aaacCTACACGGTTGATTGTAACAACGCGTTCTTCg |
| | BsaI rpsE 85-as | tatccCAACCGTGTAGCAAAAGTTGTAAAAGGTGGT |
| | BsaI rpsE 85-s | aaacACCACCTTTTACAACTTTTGCTACACGGTTGg |
| | BsaI rpsE 104-as | tatccGTAAAAGGTGGTCGTCGTTTCCGTTTCACTG |
| | BsaI rpsE 104-s | aaacCAGTGAAACGGAAACGACGACCACCTTTTACg |
| | BsaI rpsE 133-as | tatccTGCATTAGTTGTAGTTGGAGACAAAAATGGT |
| | BsaI rpsE 133-s | aaacACCATTTTTGTCTCCAACTACAACTAATGCAg |
| | BsaI rpsE 158-as | tatccAATGGTCGTGTAGGTTTCGGTACTGGTAAAG |
| | BsaI rpsE 158-s | aaacCTTTACCAGTACCGAAACCTACACGACCATTg |
| | BsaI rpsE 176-as | tatccGGTACTGGTAAAGCTCAAGAGGTACCAGAAG |
| | BsaI rpsE 176-s | aaacCTTCTGGTACCTCTTGAGCTTTACCAGTACCg |
| | BsaI rpsE 197-as | tatccGTACCAGAAGCAATCAAAAAAGCTGTTGAAG |
| | BsaI rpsE 197-s | aaacCTTCAACAGCTTTTTTGATTGCTTCTGGTACg |
| | BsaI rpsE 223-as | tatccTGAAGCAGCTAAAAAAGATTTAGTAGTTGTT |
| | BsaI rpsE 223-s | aaacAACAACTACTAAATCTTTTTTAGCTGCTTCAg |
| | BsaI rpsE 242-as | tatccTTAGTAGTTGTTCCACGTGTTGAAGGTACAA |
| | BsaI rpsE 242-s | aaacTTGTACCTTCAACACGTGGAACAACTACTAAg |
| | BsaI rpsE 262-as | tatccTGAAGGTACAACTCCACACACAATTACTGGC |
| | BsaI rpsE 262-s | aaacGCCAGTAATTGTGTGTGGAGTTGTACCTTCAg |
| | BsaI rpsE 279-as | tatccACACAATTACTGGCCGTTACGGTTCAGGAAG |
| | BsaI rpsE 279-s | aaacCTTCCTGAACCGTAACGGCCAGTAATTGTGTg |
| | BsaI rpsE 308-as | tatccAGCGTATTTATGAAACCGGCTGCACCTGGTA |
| | BsaI rpsE 308-s | aaacTACCAGGTGCAGCCGGTTTCATAAATACGCTg |
| | BsaI rpsE 329-as | tatccGCACCTGGTACAGGAGTTATCGCTGGTGGTC |
| | BsaI rpsE 329-s | aaacGACCACCAGCGATAACTCCTGTACCAGGTGCg |
| | BsaI rpsE 353-as | tatccGGTGGTCCTGTTCGTGCCGTACTTGAATTAG |
| | BsaI rpsE 353-s | aaacCTAATTCAAGTACGGCACGAACAGGACCACCg |
| | BsaI rpsE 380-as | tatccTTAGCAGGTATCACTGATATCTTAAGTAAAT |
| | BsaI rpsE 380-s | aaacATTTACTTAAGATATCAGTGATACCTGCTAAg |
| | BsaI rpsE 398-as | tatccATCTTAAGTAAATCATTAGGATCAAACACAC |
| | BsaI rpsE 398-s | aaacGTGTGTTTGATCCTAATGATTTACTTAAGATg |
| | BsaI rpsE 423-as | tatccACACACCAATCAACATGGTTCGTGCTACAAT |
| | BsaI rpsE 423-s | aaacATTGTAGCACGAACCATGTTGATTGGTGTGTg |
| | BsaI rpsE 449-as | tatccACAATCGATGGTTTACAAAACCTTAAAAATG |
| | BsaI rpsE 449-s | aaacCATTTTTAAGGTTTTGTAAACCATCGATTGTg |
| | BsaI rpsE 474-as | tatccAAAATGCTGAAGATGTTGCGAAATTACGTGG |
| | BsaI rpsE 474-s | aaacCCACGTAATTTCGCAACATCTTCAGCATTTTg |
| | BsaI ermC 33-as | tatccTGAACGAGAAAAATATAAAACACAGTCAAAA |
| | BsaI ermC 33-s | aaacTTTTGACTGTGTTTTATATTTTTCTCGTTCAg |
| | BsaI ermC 71-as | tatccACTTCAAAACATAATATAGATAAAATAATGA |
| | BsaI ermC 71-s | aaacTCATTATTTTATCTATATTATGTTTTGAAGTg |
| | BsaI ermC 102-as | tatccCAAATATAAGATTAAATGAACATGATAATAT |
| | BsaI ermC 102-s | aaacATATTATCATGTTCATTTAATCTTATATTTGg |
| | BsaI ermC 134-as | tatccTTTGAAATCGGCTCAGGAAAAGGGCATTTTA |
| | BsaI ermC 134-s | aaacTAAAATGCCCTTTTCCTGAGCCGATTTCAAAg |
| | BsaI ermC 167-as | tatccCTTGAATTAGTACAGAGGTGTAATTTCGTAA |
| | BsaI ermC 167-s | aaacTTACGAAATTACACCTCTGTACTAATTCAAGg |
| | BsaI ermC 200-as | tatccGCCATTGAAATAGACCATAAATTATGCAAAA |

| | | |
|---|---|---|
| | BsaI ermC 200-s | aaacTTTTGCATAATTTATGGTCTATTTCAATGGCg |
| | BsaI ermC 234-as | tatccCAGAAAATAAACTTGTTGATCACGATAATTT |
| | BsaI ermC 234-s | aaacAAATTATCGTGATCAACAAGTTTATTTTCTGg |
| | BsaI ermC 272-as | tatccTTAAACAAGGATATATTGCAGTTTAAATTTC |
| | BsaI ermC 272-s | aaacGAAATTTAAACTGCAATATATCCTTGTTTAAg |
| | BsaI ermC 307-as | tatccAAACCAATCCTATAAAATATTTGGTAATATA |
| | BsaI ermC 307-s | aaacTATATTACCAAATATTTTATAGGATTGGTTTg |
| | BsaI ermC 336-as | tatccTACCTTATAACATAAGTACGGATATAATACG |
| | BsaI ermC 336-s | aaacCGTATTATATCCGTACTTATGTTATAAGGTAg |
| | BsaI ermC 378-as | tatccTTGATAGTATAGCTGATGAGATTTATTTAAT |
| | BsaI ermC 378-s | aaacATTAAATAAATCTCATCAGCTATACTATCAAg |
| | BsaI ermC 418-as | tatccCGAGTTTGCTAAAAGATTATTAAATACAAAA |
| | BsaI ermC 418-s | aaacTTTTGTATTTAATAATCTTTTAGCAAACTCGg |
| | BsaI ermC 458-as | tatccGCATTATTTTTAATGGCAGAAGTTGATATTT |
| | BsaI ermC 458-s | aaacAAATATCAACTTCTGCCATTAAAAATAATGCg |
| | BsaI ermC 490-as | tatccTATATTAAGTATGGTTCCAAGAGAATATTTT |
| | BsaI ermC 490-s | aaacAAAATATTCTCTTGGAACCATACTTAATATAg |
| | BsaI ermC 522-as | tatccATCCTAAACCTAAAGTGAATAGCTCACTTAT |
| | BsaI ermC 522-s | aaacATAAGTGAGCTATTCACTTTAGGTTTAGGATg |
| | BsaI ermC 565-as | tatccAAAAAAATCAAGAATATCACACAAAGATAAA |
| | BsaI ermC 565-s | aaacTTTATCTTTGTGTGATATTCTTGATTTTTTTg |
| | BsaI ermC 600-as | tatccAGTATAATTATTTCGTTATGAAATGGGTTAA |
| | BsaI ermC 600-s | aaacTTAACCCATTTCATAACGAAATAATTATACTg |
| | BsaI ermC 639-as | tatccACAAGAAAATATTTACAAAAAATCAATTTAA |
| | BsaI ermC 639-s | aaacTTAAATTGATTTTTTGTAAATATTTTCTTGTg |
| | BsaI ermC 675-as | tatccCCTTAAAACATGCAGGAATTGACGATTTAAA |
| | BsaI ermC 675-s | aaacTTTAAATCGTCAATTCCTGCATGTTTTAAGGg |
| | BsaI ermC 714-as | tatccGCTTTGAACAATTCTTATCTCTTTTCAATAG |
| | BsaI ermC 714-s | aaacCTATTGAAAAGAGATAAGAATTGTTCAAAGCg |
| | BsaI MecA 48-as | tatccTTGTTCCACTTATTTTAATAGTTGTAGTTGT |
| | BsaI MecA 48-s | aaacACAACTACAACTATTAAAATAAGTGGAACAAg |
| | BsaI MecA 147-as | tatccATAAAAATTTCAAACAAGTTTATAAAGATAG |
| | BsaI MecA 147-s | aaacCTATCTTTATAAACTTGTTTGAAATTTTTATg |
| | BsaI MecA 247-as | tatccTTTAGGCGTTAAAGATATAAACATTCAGGAT |
| | BsaI MecA 247-s | aaacATCCTGAATGTTTATATCTTTAACGCCTAAAg |
| | BsaI MecA 337-as | tatccAACAAACTACGGTAACATTGATCGCAACGTT |
| | BsaI MecA 337-s | aaacAACGTTGCGATCAATGTTACCGTAGTTTGTTg |
| | BsaI MecA 446-as | tatccGACCAAAGCATACATATTGAAAATTTAAAAT |
| | BsaI MecA 446-s | aaacATTTTAAATTTTCAATATGTATGCTTTGGTCg |
| | BsaI MecA 554-as | tatccCCAAAGAATGTATCTAAAAAAGATTATAAAG |
| | BsaI MecA 554-s | aaacCTTTATAATCTTTTTTAGATACATTCTTTGGg |
| | BsaI MecA 648-as | tatccTACAAGATGATACCTTCGTTCCACTTAAAAC |
| | BsaI MecA 648-s | aaacGTTTTAAGTGGAACGAAGGTATCATCTTGTAg |
| | BsaI MecA 749-as | tatccAGTCGTAACTATCCTCTAGGAAAAGCGACTT |
| | BsaI MecA 749-s | aaacAAGTCGCTTTTCCTAGAGGATAGTTACGACTg |
| | BsaI MecA 849-as | tatccAAGATGATGCAGTTATTGGTAAAAAGGGACT |
| | BsaI MecA 849-s | aaacAGTCCCTTTTTACCAATAACTGCATCATCTTg |
| | BsaI MecA 963-as | tatccATACATTAATAGAGAAAAAGAAAAAAGATGG |
| | BsaI MecA 963-s | aaacCCATCTTTTTTCTTTTTCTCTATTAATGTATg |
| | BsaI MecA 1050-as | tatccTGAAAAATGATTATGGCTCAGGTACTGCTAT |
| | BsaI MecA 1050-s | aaacATAGCAGTACCTGAGCCATAATCATTTTTCAg |
| | BsaI MecA 1151-as | tatccGGCATGAGTAACGAAGAATATAATAAATTAA |
| | BsaI MecA 1151-s | aaacTTAATTTATTATATTCTTCGTTACTCATGCCg |
| | BsaI MecA 1248-as | tatccAAATATTAACAGCAATGATTGGGTTAAATAA |
| | BsaI MecA 1248-s | aaacTTATTTAACCCAATCATTGCTGTTAATATTTg |
| | BsaI MecA 1359-as | tatccTTACAAGATATGAAGTGGTAAATGGTAATAT |
| | BsaI MecA 1359-s | aaacATATTACCATTTACCACTTCATATCTTGTAAg |
| | BsaI MecA 1446-as | tatccTCGAATTAGGCAGTAAGAAATTTGAAAAAGG |
| | BsaI MecA 1446-s | aaacCCTTTTTCAAATTTCTTACTGCCTAATTCGAg |
| | BsaI MecA 1550-as | tatccAATTTAGATAATGAAATATTATTAGCTGATT |
| | BsaI MecA 1550-s | aaacAATCAGCTAATAATATTTCATTATCTAAATTg |
| | BsaI MecA 1644-as | tatccTAGAAAATAATGGCAATATTAACGCACCTCA |
| | BsaI MecA 1644-s | aaacTGAGGTGCGTTAATATTGCCATTATTTTCTAg |
| | BsaI MecA 1747-as | tatccTGATGGTATGCAACAAGTCGTAAATAAAACA |
| | BsaI MecA 1747-s | aaacTGTTTTATTTACGACTTGTTGCATACCATCAg |
| | BsaI MecA 1856-as | tatccGAAACTGGCAGACAAATTGGGTGGTTTATAT |
| | BsaI MecA 1856-s | aaacATATAAACCACCCAATTTGTCTGCCAGTTTCg |
| | BsaI MecA 1946-as | tatccAAAGGAATGGCTAGCTACAATGCCAAAATCT |
| | BsaI MecA 1946-s | aaacAGATTTTGGCATTGTAGCTAGCCATTCCTTTg |
| Construction of pKLC31 | InFpi araCosPPi NotI1-f | cttctccatgcggccatcgatgcataatgtgcctg |
| | InFpi araCosPPi NotI1-r | agcctaccggcggccTGCTGCGAGTGATTAACAAG |
| Construction of pKLC27 | InF27 araCosPPi-f | GGCAATTCCGACGTCatcgatgcataatgtgcctg |
| | InF27 araCosPPi-r | TCAAAACGCGTCGACttactgttctgtatcggctg |
| | InF27 pRC319-f | gtcgacgcgttttgaaagc |
| | InF27 pRC319-r | gacgtcggaattgccagctg |
| Construction of antibiotics | pKLC26-f | GAATTCgatcctctagagtcg |

| | | |
|---|---|---|
| resistant gene expression vector | pKLC26-r | GAGAATGGATTTTGTGATGCC |
| | HygroR-f | ACAAAATCCATTCTCatgaaaaagcctgaactcac |
| | HygroR-r | tagaggatcGAATTCttattcctttgccctcggac |
| | pAH143 GenR-f | ACAAAATCCATTCTCatgttatggagcagcaacga |
| | pAH143 GenR-r | tagaggatcGAATTCttaggtggcggtacttgggt |
| | pAH144 SpcR-StrR-f | ACAAAATCCATTCTCatgcgctcacgcaactggtc |
| | pAH144 SpcR-StrR-r | tagaggatcGAATTCttatttgccgactaccttgg |
| | pAH145 TmpR-f | ACAAAATCCATTCTCatgggtcaaagtagcgatga |
| | pAH145 TmpR-r | tagaggatcGAATTCttaggccacacgttcaagtg |
| | pRTKH2 ErmR-f | ACAAAATCCATTCTCatgaacaaaaatataaaatattctc |
| | pRTKH2 ErmR-r | tagaggatcGAATTCttatttcctcccgttaaataatag |
| | PuroR-f | ACAAAATCCATTCTCggtgcgcctcgccacccg |
| | PuroR-r | tagaggatcGAATTCtcaggcaccgggcttgcg |
| Construction of pKLC44 | InF44 pKLC42-f | cagggcgcgtctcgaGtctagaCGATCGCTTGATAAGG |
| | InF44 pKLC42-r | acaggagtccaagcgAGCTcgaaagcaaattcgCTCGAG |
| Genetic modification of E. coli | K12 genome-in pKLC26-s | cccttcaaccttagcagtagcgtgggatgatttcacaattagaaagacctTGACGCACACCGTGGAAAC |
| | K12 genome-in pKLC26Cm-as | tgtcctgcacgacgccttgcgtcactagcctcttctctagctcatcatgcTGAGACGTTGATCGGCACG |
| Detection primers for S. aureus | SA 23S rRNA det-s | AAGCGAGTCTGAATAGGGCG |
| | SA 23S rRNA det-as | AACGTAAGTCGGTTCGGTCC |
| | SA RPF3757 MecA det-s | TGGCTCAGGTACTGCTATCC |
| | SA RPF3757 MecA det-as | AGACGTCATATGAAGGTGTGC |
| | SA RPF3757 ermC det-s | CTTTGAAATCGGCTCAGGAAAAGG |
| | SA RPF3757 ermC det-as | GGCAGTTACGAAATTACACCTCTG |
| | SA RPF3757 rpsE det-s | GGTCGTCGTTTCCGTTTCAC |
| | SA RPF3757 rpsE det-as | TGCTTCTGGTACCTCTTGAGC |
| Construction of target expression vectors for S. aureus | USA300p3 ermC clo BamHI-s | ATATGGATCCATGAACGAGAAAAATATAAAACACAG |
| | USA300p3 ermC clo EcoRI-as | ATATGAATTCCAAAAGACATAATCGATTCAC |
| | rpsE clo BamHI-s | atatGGATCCatggctcgtagagaagaag |
| | rpsE clo SacI-as | atatGAGCTCcctccttaattgtataattcttctac |
| | USA300g MecA clo BamHI-s2 | ATATGGATCCGTAGTCTTATATAAGGAGTATATTG |
| | USA300g MecA clo EcoRI-as2 | ATATGAATTCTAAGGGAGAAGTAACAGCAC |
| | RFP clo BamHI-s | ATATGGATCCATGGCGAGTAGCGAAGAC |
| | RFP clo EcoRI-as | ATATGAATTCTTAAGCACCGGTGGAGTG |
| Construction of stx1/2 expression vector | pKLC26 stx1 partial PCR SacI-as | GTGGAGCTCGGTCATGGCATTTCCACTAAACTCCATTAAGAGAATGGATTTTGTGATGC |
| | pKLC26 stx1 partial PCR SacI -s | ACCGAGCTCCACCGGAAGAAGTGGAACTCACACTGAACGAATTCgatcctctagagtc |
| | pKLC26 stx2 partial PCR SacI -as | GCGGAGCTCGGTCATTGTATTACCACTGAACTCCATTAAGAGAATGGATTTTGTGATGC |
| | pKLC26 stx2 partial PCR SacI -s | ACCGAGCTCCGCCGGGAGACGTGGACCTCACTCTGAACGAATTCgatcctctagagtc |
| Construction of SaPI-based CapsidCas13a plasmid | BAPup 7272-F | AATTCCTGCAGCCCGTTGACGAGGTTGGTAATGGCAC |
| | BAPup 8172-R | CCCTGTTGATACCGGGATACATCACTTGTTTTGCCGTC |
| | BAPdown 12224-F | TCGGCGCAAAGTGCGGGGCTCTCCACTTACAAAGGT |
| | BAPdown 13089-R | TTTGCCGTTACGCACGGGCACTCTTTTGTGTATAACCG |
| | LsC2c2 mecA5-F | CCGGTATCAACAGGGATGCCTACAGCATCCAGGGT |
| | LsC2c2 mecA5-R | CGCACTTTGCGCCGAGAACCTTCGAAAAACCGCCC |
| | LsC2c2 188-R | CGCACTTTGCGCCGACCCATCCTAATTTCTTGCTGATGAG |
| Construction of pIMAY mecA deletion construct | USA300-C02 906894-F | AATTCCTGCAGCCCGCGCCCAAAGCTTCTTTAGCTG |
| | USA300-C02 907924-R | CCCTGTTGATACCGGTGTGATATGGAGGTGTAGAAGGTG |
| | USA300-C02 910027-F | TCGGCGCAAAGTGCGCCGTAACGATGGTTGCTTCAC |
| | USA300-C02 910980-R | TTTGCCGTTACGCACCTGGACCGAATGGACTAGCA |

**Table S4. List of depletion efficiency of sequence reads**

| Position in IMP-1 | Spacer sequence | Depletion efficiency | Survival rate of *E. coli* IMP-1 | Sequence reads from *E. coli* mock | | | | Sequence reads from *E. coli* IMP-1 | | | |
|---|---|---|---|---|---|---|---|---|---|---|---|
| | | | | Rep.1 | Rep.2 | Rep.3 | Rep.4 | Rep.1 | Rep.2 | Rep.3 | Rep.4 |
| IMP-1_563 | GACTTTGGCCAAGCTTCTATATTTGCGT | 364.1 | 0.0027 | 56 | 75 | 47 | 67 | 1 | 0 | 3 | 0 |
| IMP-1_566 | GCGGACTTTGGCCAAGCTTCTATATTTG | 256.8 | 0.0039 | 49 | 70 | 40 | 57 | 1 | 1 | 3 | 0 |
| IMP-1_702 | TTTTGATGGTTTTTTACTTTCGTTTAAC | 236.9 | 0.0042 | 40 | 106 | 65 | 68 | 0 | 2 | 5 | 0 |
| IMP-1_562 | ACTTTGGCCAAGCTTCTATATTTGCGTC | 220.8 | 0.0045 | 44 | 88 | 58 | 70 | 0 | 4 | 3 | 0 |
| IMP-1_370 | TGGCTTGAACCTTACCGTCTTTTTTAAG | 201.4 | 0.0050 | 53 | 89 | 72 | 57 | 0 | 1 | 7 | 0 |
| IMP-1_565 | CGGACTTTGGCCAAGCTTCTATATTTGC | 175.4 | 0.0057 | 55 | 80 | 55 | 46 | 2 | 0 | 3 | 3 |
| IMP-1_193 | TATCTTTAGCCGTAAATGGAGTGTCAAT | 144.6 | 0.0069 | 49 | 65 | 49 | 56 | 1 | 1 | 3 | 4 |
| IMP-1_274 | TGCTGTCGCTATGAAAATGAGAGGAAAT | 144.6 | 0.0069 | 42 | 52 | 58 | 67 | 0 | 1 | 6 | 2 |
| IMP-1_648 | TTTCAAGAGTGATGCGTCTCCAACTTCA | 136.2 | 0.0073 | 47 | 91 | 60 | 54 | 0 | 2 | 6 | 3 |
| IMP-1_11 | CAAAACAAAAATATAAAGAATACAGAT | 128.8 | 0.0078 | 40 | 66 | 46 | 43 | 1 | 1 | 5 | 2 |
| IMP-1_609 | AACAACCAGTTTTGCCTTACCATATTTG | 123.2 | 0.0081 | 41 | 67 | 46 | 74 | 4 | 1 | 6 | 0 |
| IMP-1_703 | GTTTTGATGGTTTTTTACTTTCGTTTAA | 120.4 | 0.0083 | 54 | 69 | 69 | 51 | 2 | 2 | 3 | 5 |
| IMP-1_208 | TGACTAACTTTTCAGTATCTTTAGCCGT | 106.5 | 0.0094 | 46 | 65 | 55 | 49 | 0 | 3 | 4 | 5 |
| IMP-1_322 | ATGCATACGTGGGGATAGATCGAGAATT | 99.8 | 0.0100 | 49 | 75 | 56 | 55 | 3 | 1 | 6 | 4 |
| IMP-1_206 | ACTAACTTTTCAGTATCTTTAGCCGTAA | 99.7 | 0.0100 | 48 | 67 | 54 | 49 | 5 | 0 | 7 | 1 |
| IMP-1_339 | TTCATTTGTTAATTCAGATGCATACGTG | 93.1 | 0.0107 | 30 | 103 | 50 | 52 | 1 | 4 | 6 | 4 |
| IMP-1_181 | TAAATGGAGTGTCAATTAGGTAAGCCTC | 92.4 | 0.0108 | 35 | 41 | 49 | 46 | 5 | 0 | 5 | 1 |
| IMP-1_637 | ATGCGTCTCCAACTTCACTGTGACTTGG | 88.6 | 0.0113 | 39 | 36 | 36 | 38 | 2 | 3 | 4 | 1 |
| IMP-1_368 | GCTTGAACCTTACCGTCTTTTTTAAGCA | 87.1 | 0.0115 | 62 | 77 | 56 | 54 | 3 | 6 | 6 | 2 |
| IMP-1_652 | TAAGTTTCAAGAGTGATGCGTCTCCAAC | 86.4 | 0.0116 | 43 | 62 | 56 | 57 | 6 | 2 | 6 | 1 |
| IMP-1_269 | TCGCTATGAAAATGAGAGGAAATGCTGC | 81.0 | 0.0123 | 39 | 22 | 23 | 25 | 4 | 0 | 4 | 0 |
| IMP-1_272 | CTGTCGCTATGAAAATGAGAGGAAATGC | 72.1 | 0.0139 | 38 | 75 | 41 | 40 | 5 | 3 | 3 | 5 |
| IMP-1_376 | AATTTGTGGCTTGAACCTTACCGTCTTT | 70.2 | 0.0142 | 40 | 93 | 59 | 56 | 9 | 3 | 6 | 3 |
| IMP-1_266 | CTATGAAAATGAGAGGAAATGCTGCCTT | 68.8 | 0.0145 | 8 | 31 | 10 | 32 | 1 | 2 | 3 | 1 |
| IMP-1_120 | AACAACGCCCCACCCGTTAACTTCTTCA | 65.4 | 0.0153 | 43 | 70 | 54 | 64 | 10 | 3 | 8 | 0 |
| IMP-1_363 | AACCTTACCGTCTTTTTTAAGCAGTTCA | 65.1 | 0.0154 | 37 | 81 | 68 | 55 | 2 | 8 | 12 | 0 |
| IMP-1_23 | GTAGCAATGCTGCAAAACAAAAATATAA | 64.8 | 0.0154 | 47 | 57 | 61 | 42 | 3 | 7 | 4 | 5 |
| IMP-1_392 | TTAACTCCGCTAAATGAATTTGTGGCTT | 64.4 | 0.0155 | 11 | 25 | 7 | 22 | 0 | 0 | 3 | 3 |
| IMP-1_471 | CAACCAAACCACTACGTTATCTGGAGTG | 62.0 | 0.0161 | 42 | 73 | 50 | 54 | 6 | 9 | 4 | 2 |
| IMP-1_615 | ACTTGGAACAACCAGTTTTGCCTTACCA | 61.9 | 0.0162 | 28 | 39 | 26 | 32 | 3 | 3 | 5 | 1 |
| IMP-1_170 | TCAATTAGGTAAGCCTCAGCATTTACAA | 60.8 | 0.0164 | 11 | 84 | 26 | 53 | 0 | 12 | 3 | 2 |
| IMP-1_104 | TTAACTTCTTCAAACGAAGTATGAACAT | 57.1 | 0.0175 | 17 | 58 | 29 | 40 | 3 | 6 | 3 | 3 |
| IMP-1_472 | GCAACCAAACCACTACGTTATCTGGAGT | 54.9 | 0.0182 | 44 | 70 | 46 | 80 | 2 | 8 | 10 | 6 |
| IMP-1_121 | GAACAACGCCCCACCCGTTAACTTCTTC | 50.7 | 0.0197 | 39 | 65 | 43 | 49 | 5 | 14 | 4 | 0 |
| IMP-1_639 | TGATGCGTCTCCAACTTCACTGTGACTT | 49.3 | 0.0203 | 26 | 31 | 33 | 26 | 6 | 2 | 4 | 2 |
| IMP-1_342 | CAGTTCATTTGTTAATTCAGATGCATAC | 49.2 | 0.0203 | 12 | 88 | 31 | 51 | 1 | 13 | 4 | 4 |
| IMP-1_448 | GAGTGTGTCCCGGGCCTGGATAAAAAAC | 48.3 | 0.0207 | 50 | 80 | 51 | 63 | 4 | 11 | 13 | 2 |
| IMP-1_555 | CCAAGCTTCTATATTTGCGTCACCCAAA | 46.1 | 0.0217 | 45 | 78 | 53 | 49 | 12 | 5 | 7 | 5 |
| IMP-1_550 | CTTCTATATTTGCGTCACCCAAATTGCC | 43.0 | 0.0233 | 57 | 72 | 46 | 49 | 7 | 6 | 13 | 5 |
| IMP-1_610 | GAACAACCAGTTTTGCCTTACCATATTT | 42.2 | 0.0237 | 35 | 62 | 61 | 41 | 9 | 10 | 7 | 2 |
| IMP-1_81 | AACATAAACGCCTTCATCAAGCTTTTCA | 42.2 | 0.0237 | 49 | 76 | 57 | 52 | 5 | 10 | 14 | 4 |
| IMP-1_28 | CTGCGGTAGCAATGCTGCAAAACAAAAA | 38.4 | 0.0260 | 53 | 104 | 64 | 57 | 11 | 6 | 15 | 11 |
| IMP-1_367 | CTTGAACCTTACCGTCTTTTTTAAGCAG | 36.5 | 0.0274 | 54 | 94 | 41 | 81 | 10 | 16 | 11 | 7 |
| IMP-1_600 | TTTTGCCTTACCATATTTGGACTTTAAT | 35.5 | 0.0282 | 51 | 66 | 38 | 54 | 12 | 8 | 9 | 6 |
| IMP-1_88 | AAGTATGAACATAAACGCCTTCATCAAG | 30.3 | 0.0330 | 54 | 70 | 49 | 51 | 7 | 13 | 17 | 7 |
| IMP-1_660 | CTCTAATGTAAGTTTCAAGAGTGATGCG | 30.3 | 0.0330 | 45 | 71 | 61 | 47 | 10 | 15 | 13 | 6 |
| IMP-1_126 | TTTAGGAACAACGCCCCACCCGTTAACT | 30.2 | 0.0331 | 39 | 62 | 62 | 71 | 16 | 9 | 16 | 5 |
| IMP-1_697 | ATGGTTTTTTACTTTCGTTTAACCCTTT | 26.9 | 0.0372 | 30 | 59 | 46 | 46 | 8 | 13 | 9 | 10 |
| IMP-1_60 | CTTTTCAATTTTTAAATCTGGCAAAGAC | 26.2 | 0.0382 | 35 | 83 | 39 | 41 | 10 | 13 | 10 | 12 |
| IMP-1_383 | CTAAATGAATTTGTGGCTTGAACCTTAC | 23.4 | 0.0428 | 38 | 72 | 79 | 51 | 24 | 15 | 15 | 7 |
| IMP-1_227 | CCACGCTCCACAAACCAAGTGACTAACT | 22.7 | 0.0441 | 37 | 86 | 58 | 44 | 22 | 13 | 19 | 5 |
| IMP-1_141 | AACCACCAAACCATGTTTAGGAACAACC | 22.6 | 0.0443 | 31 | 54 | 35 | 51 | 11 | 16 | 11 | 7 |
| IMP-1_250 | AAATGCTGCCTTTTATTTTATAGCCACG | 22.2 | 0.0450 | 13 | 24 | 19 | 15 | 2 | 6 | 9 | 2 |
| IMP-1_162 | GTAAGCCTCAGCATTTACAAGAACCACC | 21.7 | 0.0460 | 54 | 53 | 52 | 42 | 20 | 15 | 20 | 0 |
| IMP-1_261 | AAAATGAGAGGAAATGCTGCCTTTTATT | 21.5 | 0.0464 | 3 | 53 | 6 | 25 | 2 | 10 | 3 | 9 |
| IMP-1_556 | GCCAAGCTTCTATATTTGCGTCACCCAA | 21.4 | 0.0467 | 46 | 63 | 37 | 52 | 8 | 17 | 18 | 12 |
| IMP-1_184 | CCGTAAATGGAGTGTCAATTAGGTAAGC | 21.0 | 0.0476 | 50 | 63 | 32 | 46 | 8 | 12 | 23 | 11 |
| IMP-1_710 | TTGCTTGGTTTTGATGGTTTTTTACTTT | 20.8 | 0.0482 | 34 | 68 | 41 | 49 | 15 | 18 | 8 | 14 |
| IMP-1_40 | GCAAAGACTCTGCTGCGGTAGCAATGCT | 20.2 | 0.0495 | 3 | 81 | 22 | 57 | 13 | 12 | 11 | 12 |
| IMP-1_157 | CCTCAGCATTTACAAGAACCACCAAACC | 19.9 | 0.0503 | 56 | 31 | 71 | 16 | 30 | 0 | 22 | 0 |
| IMP-1_85 | TATGAACATAAACGCCTTCATCAAGCTT | 19.4 | 0.0515 | 20 | 67 | 26 | 47 | 6 | 22 | 4 | 17 |
| IMP-1_680 | TTTAACCCTTTAACCGCCTGCTCTAATG | 18.6 | 0.0538 | 43 | 97 | 40 | 64 | 28 | 20 | 21 | 9 |
| IMP-1_617 | TGACTTGGAACAACCAGTTTTGCCTTAC | 17.9 | 0.0558 | 37 | 69 | 36 | 60 | 14 | 21 | 12 | 20 |
| IMP-1_632 | TCTCCAACTTCACTGTGACTTGGAACAA | 17.5 | 0.0573 | 42 | 82 | 48 | 60 | 26 | 23 | 16 | 14 |
| IMP-1_447 | AGTGTGTCCCGGGCCTGGATAAAAAACT | 17.0 | 0.0590 | 44 | 109 | 67 | 71 | 31 | 30 | 20 | 21 |
| IMP-1_19 | CAATGCTGCAAAACAAAAATATAAAGAA | 16.6 | 0.0601 | 55 | 74 | 59 | 75 | 28 | 34 | 21 | 11 |
| IMP-1_161 | TAAGCCTCAGCATTTACAAGAACCACCA | 16.4 | 0.0610 | 30 | 69 | 42 | 52 | 24 | 21 | 14 | 11 |
| IMP-1_353 | TCTTTTTTAAGCAGTTCATTTGTTAATT | 16.1 | 0.0621 | 35 | 71 | 52 | 48 | 20 | 21 | 17 | 18 |
| IMP-1_34 | ACTCTGCTGCGGTAGCAATGCTGCAAAA | 15.4 | 0.0650 | 51 | 87 | 57 | 56 | 30 | 26 | 28 | 13 |
| IMP-1_536 | TCACCCAAATTGCCTAAACCGTACGGTT | 14.4 | 0.0697 | 31 | 63 | 33 | 42 | 15 | 26 | 13 | 16 |

| | | | | | | | | | | | |
|---|---|---|---|---|---|---|---|---|---|---|---|
| IMP-1_291 | CCACTCTATTCCGCCCGTGCTGTCGCTA | 13.9 | 0.0721 | 4 | 23 | 9 | 13 | 5 | 7 | 7 | 2 |
| IMP-1_251 | GAAATGCTGCCTTTTATTTTATAGCCAC | 13.2 | 0.0759 | 11 | 19 | 4 | 17 | 4 | 8 | 3 | 8 |
| IMP-1_445 | TGTGTCCCGGGCCTGGATAAAAAACTTC | 12.1 | 0.0824 | 16 | 93 | 23 | 62 | 7 | 58 | 4 | 26 |
| IMP-1_664 | CCTGCTCTAATGTAAGTTTCAAGAGTGA | 12.1 | 0.0829 | 39 | 75 | 53 | 42 | 33 | 25 | 31 | 14 |
| IMP-1_39 | CAAAGACTCTGCTGCGGTAGCAATGCTG | 12.0 | 0.0831 | 29 | 58 | 37 | 38 | 29 | 24 | 18 | 9 |
| IMP-1_174 | AGTGTCAATTAGGTAAGCCTCAGCATTT | 10.8 | 0.0924 | 36 | 71 | 44 | 51 | 38 | 35 | 18 | 20 |
| IMP-1_22 | TAGCAATGCTGCAAAACAAAAATATAAA | 10.7 | 0.0937 | 51 | 71 | 61 | 45 | 27 | 45 | 32 | 23 |
| IMP-1_580 | ACTTTAATAATTTGGCGGACTTTGGCCA | 10.6 | 0.0945 | 41 | 71 | 62 | 68 | 30 | 43 | 39 | 24 |
| IMP-1_25 | CGGTAGCAATGCTGCAAAACAAAAATAT | 10.0 | 0.1002 | 28 | 71 | 33 | 41 | 30 | 30 | 20 | 23 |
| IMP-1_430 | GATAAAAAACTTCAATTTTATTTTTAAC | 9.8 | 0.1017 | 30 | 83 | 67 | 50 | 47 | 36 | 35 | 21 |
| IMP-1_399 | CCAATAGTTAACTCCGCTAAATGAATTT | 8.1 | 0.1230 | 38 | 36 | 54 | 17 | 36 | 20 | 31 | 19 |
| IMP-1_511 | GTTTAATAAAACAACCACCGAATAATAT | 7.4 | 0.1344 | 39 | 55 | 22 | 43 | 22 | 58 | 27 | 20 |
| IMP-1_222 | CTCCACAAACCAAGTGACTAACTTTTCA | 6.2 | 0.1612 | 53 | 85 | 78 | 46 | 79 | 89 | 50 | 33 |
| IMP-1_641 | AGTGATGCGTCTCCAACTTCACTGTGAC | 6.2 | 0.1612 | 18 | 24 | 18 | 12 | 19 | 15 | 25 | 10 |
| IMP-1_253 | AGGAAATGCTGCCTTTTATTTTATAGCC | 5.9 | 0.1682 | 0 | 6 | 4 | 5 | 4 | 1 | 3 | 7 |
| IMP-1_172 | TGTCAATTAGGTAAGCCTCAGCATTTAC | 5.7 | 0.1765 | 44 | 90 | 44 | 47 | 58 | 81 | 44 | 53 |
| IMP-1_595 | CCTTACCATATTTGGACTTTAATAATTT | 5.3 | 0.1879 | 42 | 74 | 53 | 45 | 49 | 86 | 57 | 47 |
| IMP-1_175 | GAGTGTCAATTAGGTAAGCCTCAGCATT | 5.2 | 0.1916 | 23 | 30 | 32 | 23 | 46 | 29 | 27 | 21 |
| IMP-1_310 | GGATAGATCGAGAATTAAGCCACTCTAT | 5.2 | 0.1924 | 34 | 74 | 47 | 47 | 53 | 78 | 53 | 47 |
| IMP-1_151 | CATTTACAAGAACCACCAAACCATGTTT | 5.1 | 0.1944 | 44 | 63 | 45 | 54 | 72 | 77 | 52 | 37 |
| IMP-1_436 | GGCCTGGATAAAAAACTTCAATTTTATT | 5.1 | 0.1963 | 30 | 42 | 29 | 19 | 38 | 46 | 25 | 31 |
| IMP-1_429 | ATAAAAAACTTCAATTTTATTTTTAACT | 4.7 | 0.2109 | 17 | 48 | 33 | 32 | 28 | 62 | 39 | 34 |
| IMP-1_309 | GATAGATCGAGAATTAAGCCACTCTATT | 4.6 | 0.2164 | 44 | 63 | 51 | 34 | 61 | 74 | 63 | 49 |
| IMP-1_538 | CGTCACCCAAATTGCCTAAACCGTACGG | 4.4 | 0.2276 | 44 | 76 | 52 | 52 | 68 | 107 | 53 | 75 |
| IMP-1_707 | CTTGGTTTTGATGGTTTTTTACTTTCGT | 4.3 | 0.2353 | 52 | 83 | 54 | 67 | 110 | 101 | 76 | 71 |
| IMP-1_435 | GCCTGGATAAAAAACTTCAATTTTATTT | 4.1 | 0.2457 | 36 | 51 | 48 | 41 | 78 | 81 | 53 | 45 |
| IMP-1_311 | GGGATAGATCGAGAATTAAGCCACTCTA | 3.9 | 0.2536 | 43 | 59 | 61 | 42 | 76 | 87 | 67 | 79 |
| IMP-1_113 | CCCCACCCGTTAACTTCTTCAAACGAAG | 3.9 | 0.2561 | 42 | 72 | 52 | 56 | 100 | 109 | 57 | 72 |
| IMP-1_298 | AATTAAGCCACTCTATTCCGCCCGTGCT | 3.9 | 0.2566 | 13 | 17 | 13 | 16 | 23 | 28 | 20 | 19 |
| IMP-1_308 | ATAGATCGAGAATTAAGCCACTCTATTC | 3.5 | 0.2875 | 35 | 52 | 47 | 55 | 83 | 120 | 57 | 63 |
| IMP-1_300 | AGAATTAAGCCACTCTATTCCGCCCGTG | 3.4 | 0.2967 | 15 | 26 | 24 | 28 | 41 | 63 | 39 | 21 |
| IMP-1_245 | CTGCCTTTTATTTTATAGCCACGCTCCA | 3.3 | 0.3038 | 11 | 43 | 24 | 25 | 52 | 64 | 26 | 44 |
| IMP-1_510 | TTTAATAAAACAACCACCGAATAATATT | 3.2 | 0.3084 | 44 | 83 | 51 | 50 | 101 | 151 | 85 | 81 |
| IMP-1_634 | CGTCTCCAACTTCACTGTGACTTGGAAC | 3.2 | 0.3108 | 41 | 64 | 44 | 41 | 93 | 110 | 80 | 68 |
| IMP-1_304 | ATCGAGAATTAAGCCACTCTATTCCGCC | 3.1 | 0.3204 | 32 | 62 | 33 | 51 | 75 | 122 | 73 | 69 |
| IMP-1_581 | GACTTTAATAATTTGGCGGACTTTGGCC | 3.1 | 0.3243 | 42 | 96 | 46 | 92 | 146 | 198 | 94 | 94 |
| IMP-1_255 | AGAGGAAATGCTGCCTTTTATTTTATAG | 3.1 | 0.3244 | 6 | 2 | 6 | 0 | 13 | 2 | 6 | 6 |
| IMP-1_693 | TTTTTTACTTTCGTTTAACCCTTTAACC | 2.9 | 0.3508 | 53 | 52 | 51 | 44 | 115 | 137 | 93 | 72 |
| IMP-1_278 | CCCGTGCTGTCGCTATGAAAATGAGAGG | 2.7 | 0.3765 | 42 | 81 | 33 | 50 | 104 | 167 | 94 | 96 |
| IMP-1_331 | TTAATTCAGATGCATACGTGGGGATAGA | 2.5 | 0.4003 | 28 | 46 | 16 | 18 | 86 | 80 | 41 | 50 |
| IMP-1_694 | GTTTTTTACTTTCGTTTAACCCTTTAAC | 2.4 | 0.4211 | 30 | 54 | 38 | 49 | 113 | 147 | 82 | 86 |
| IMP-1_455 | TTATCTGGAGTGTGTCCCGGGCCTGGAT | 2.2 | 0.4635 | 40 | 44 | 63 | 41 | 109 | 178 | 109 | 122 |
| IMP-1_443 | TGTCCCGGGCCTGGATAAAAAACTTCAA | 2.0 | 0.5067 | 63 | 72 | 63 | 55 | 181 | 249 | 144 | 188 |
| IMP-1_242 | CCTTTTATTTTATAGCCACGCTCCACAA | 2.0 | 0.5086 | 15 | 26 | 23 | 23 | 78 | 88 | 57 | 40 |
| IMP-1_491 | AATAATATTTTCCTTTCAGGCAACCAAA | 1.8 | 0.5708 | 44 | 27 | 53 | 21 | 217 | 71 | 161 | 43 |
| IMP-1_441 | TCCCGGGCCTGGATAAAAAACTTCAATT | 1.7 | 0.5785 | 39 | 81 | 29 | 56 | 152 | 263 | 131 | 159 |
| IMP-1_14 | CTGCAAAACAAAAATATAAAGAATACAC | 1.7 | 0.5972 | 17 | 44 | 24 | 35 | 55 | 205 | 54 | 112 |
| IMP-1_515 | TACGGTTTAATAAAACAACCACCGAATA | 1.6 | 0.6203 | 48 | 92 | 67 | 68 | 260 | 338 | 222 | 194 |
| IMP-1_434 | CCTGGATAAAAAACTTCAATTTTATTTT | 1.2 | 0.8093 | 26 | 84 | 55 | 46 | 243 | 338 | 209 | 225 |
| IMP-1_524 | CCTAAACCGTACGGTTTAATAAAACAAC | 1.1 | 0.8718 | 50 | 71 | 59 | 56 | 295 | 455 | 210 | 263 |
| IMP-1_71 | CCTTCATCAAGCTTTTCAATTTTTAAAT | 1.0 | 1.0207 | 46 | 58 | 51 | 39 | 321 | 364 | 252 | 240 |
| NC | GGAGACCGAGATTGGTCTC | 1.0 | 1.0000 | 45 | 92 | 58 | 74 | 380 | 574 | 254 | 391 |
